# scREAD: A single-cell RNA-Seq database for Alzheimer’s Disease

**DOI:** 10.1101/2020.08.06.240044

**Authors:** Jing Jiang, Cankun Wang, Ren Qi, Hongjun Fu, Qin Ma

**Affiliations:** Department of Biomedical Informatics, The Ohio State University, OH, USA; Department of Neuroscience, The Ohio State University, OH, USA

## Abstract

**Summary:** Alzheimer’s disease (AD) is a progressive neurodegenerative disorder of the brain and the most common form of dementia among the elderly. The single-cell RNA-sequencing (scRNA-Seq) and single-nucleus RNA-sequencing (snRNA-Seq) techniques are extremely useful for dissecting the function/dysfunction of highly heterogeneous cells in the brain at the single-cell level, and the corresponding data analyses can significantly improve our understanding of why particular cells are vulnerable in AD. We developed an integrated database named scREAD (<u>s</u>ingle-<u>c</u>ell <u>R</u>NA-Seq databas<u>e</u> for <u>A</u>lzheimer’s <u>D</u>isease), which is the first database dedicated to the management of all the existing scRNA-Seq and snRNA-Seq datasets from human postmortem brain tissue with AD and mouse models with AD pathology. scREAD provides comprehensive analysis results for 55 datasets from eight brain regions, including control atlas construction, cell type prediction, identification of differentially expressed genes, and identification of cell-type-specific regulons.

**Availability and Implementation:** scREAD is a one-stop and user-friendly interface and freely available at https://bmbls.bmi.osumc.edu/scread/. The backend workflow can be downloaded from https://github.com/OSU-BMBL/scread/tree/master/script, to enable more discovery-driven analyses.

**Supplementary information:** Supplementary data are available at *Bioinformatics* online.

## 1 Introduction

Alzheimer’s disease (AD) is the most common cause of dementia. Currently, there are an estimated 5.8 million Americans age 65 or older suffering from AD (Claxton, et al., 2015). Only after years of brain changes do individuals experience noticeable symptoms, such as memory loss and language problems (Shinagawa, 2016). Symptoms occur because neurons in parts of the brain involved in thinking, learning, and memory have been damaged or destroyed, probably by the accumulation of amyloid beta (Aβ) protein and tau protein aggregates and the neuroinflammation (Dolgin, 2018; Mucke, 2009). Unfortunately, there is no effective therapeutics that can cure or alter the disease process (Gao, et al., 2016). Furthermore, molecular mechanisms underlying AD, especially the cellular vulnerability, are poorly understood.

Previous studies have demonstrated that AD pathology differs in age, gender, brain regions and cell types (Ewers, et al., 2011; Mucke, 2009; Sala Frigerio, et al., 2019). In order to study the cellular heterogeneity of the brain and reveal the complex cellular changes in the AD brain by profiling tens of thousands of individual cells, single-cell RNA sequencing (scRNA-seq) provides an alternative method (Mathys, et al., 2017). The scRNA-Seq can reveal complex and rare cell populations, uncover regulatory relationships between genes, and track the trajectories of distinct cell lineages in development (Grubman, et al., 2019). The single-cell view of AD pathology paints a unique cellular-level view of transcriptional alterations associated with AD pathology and significantly improves our understanding of the pathogenesis of AD (Del-Aguila, et al., 2019; Mathys, et al., 2019).

Here we developed a novel database called scREAD (<u>s</u>ingle-<u>c</u>ell <u>R</u>NA-<u>S</u>eq database for <u>A</u>lzheimer’s <u>D</u>isease), which provides comprehensive analysis results of all the existing scRNA-Seq and snRNA-Seq datasets collected from GEO (Barrett, et al., 2013) and Synapse databases. The scREAD has several key features, namely, (i) it is the first-of-its-kind database with a collection of all the 12 existing human and mouse AD scRNA-Seq and snRNA-Seq datasets from the public domain (Supplementary Table S1); (ii) it re-defines the 12 datasets into 55 datasets, each of which corresponds to a specific species (human or mouse), gender (male or female), brain region (entorhinal cortex, prefrontal cortex, superior frontal gyrus, cortex, cerebellum, cerebral cortex, or hippocampus) (Supplementary Table S2), disease or control, and age stage (7 months, 15 months, or 20 months for mice, and 50-90+ years old for human) (Supplementary Table S3); (iii) it provides comprehensive analysis results for each of the 55 datasets, including but not limited to the construction of control atlas, cell clustering, prediction of cell types, identification of differentially expressed genes (DEGs), and identification of cell-type-specific regulons (CTSRs) in support of the in-depth analysis of heterogeneous regulatory mechanisms; (iv) all these analysis results are visualized through a one-stop and user-friendly interface to free AD biologists from programming burdens (Supplementary Tutorial part 1-5); (v) the backend workflow enabling all the above computational analyses is freely accessable as stand-alone one-line-command scripts in R (Supplementary Tutorial part 6).

## 2 Functions and implementation

There are four major functionalities in scREAD (*i*) Construction of control atlas for different human and mouse brain regions; (*ii*) Identification of human and mouse disease cell types based on the control atlas and subclusters; (*iii*) Identification of DEGs for each cell type among different conditions and functional enrichment analysis of DEGs; and (*iv*) Identification of CTSRs. The description of the pipeline (Fig. 1A) and these four functions are described below.

**Fig. 1.**
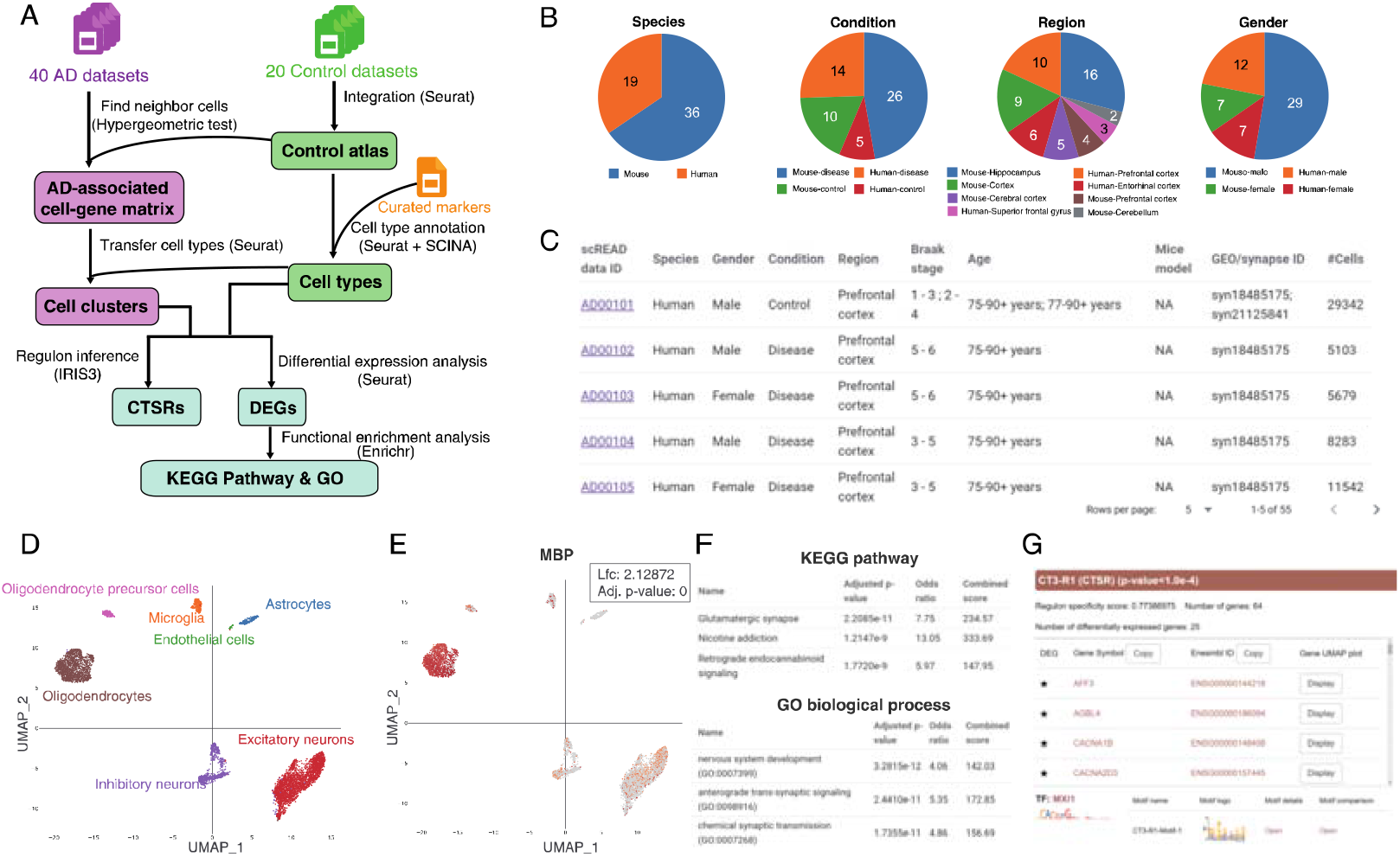
scREAD overview. (**A**) Backend workflow of scREAD; (**B**) General statistical distribution of all the 55 datasets; (**C**) General information table on the homepage; (**D**) UMAP plot colored by cell type on an AD disease dataset (AD00103); (**E**) UMAP plot of expression distribution of oligodendrocyte marker gene MBP in the same dataset; (**F**) The functional enrichment analysis of selected DEGs; (**G**) The details of the top CTSR of cell type three.

### i. Construction of control atlas for different human and mouse brain regions

We constructed 15 control cell atlases of human and mouse based on 55 scRNA-Seq and snRNA-Seq datasets, which cover eight brain regions, two genders, and different mouse and human ages, totally 579,392 cells (Fig. 1B, C). Cell types of these 15 control atlases were assigned using Seurat (Stuart, et al., 2019) and SCINA (Zhang, et al., 2019), and the known cell type markers genes used in this process were collected from literature and PanglaoDB (Franzen, et al., 2019) (Supplementary Table S4).

### ii. Identification of human and mouse disease cell types based on the control atlas and subclusters

Not all cells collected from AD patient samples are malignant, and normal control cells are included. These control cells maintain distinct regulatory mechanisms and gene expression patterns compared to AD cells and will disturb the accurate identification of AD cell types. Thus, we removed these control cells from AD data to identify real AD-associated cells. Using the human and mouse control atlas, we sought to project AD-associated cells onto the control atlas at single-cell resolution to identify human and mouse disease cell types (Fig. 1D). For each cell type, we have carried out the subcluster finding analysis for investigating subcluster-specific changes and functional diversity occurring in AD.

### iii. Identification of DEGs for each cell type among different conditions and functional enrichment analysis of DEGs

Differential gene expression analysis was performed using Seurat (Fig. 1E), including cell-type-specific genes, subcluster-specific genes, and cell-type-pairwise DEGs within one dataset or between datasets based on diverse conditions (Supplementary Table S5). Enrichr (Kuleshov, et al., 2016) was used to perform the enrichment analysis of the DEGs against different functional annotation databases (Fig. 1F).

### iv. Identification of CTSRs

The CTSRs were identified using the IRIS3 (Ma, et al., 2020), which is an integrated web server for CTSR inference from Human or Mouse Single-cell RNA-Seq data (Fig. 1G). Each regulon was named by the cell type index and regulon number and was ranked in decreasing order of regulon specificity score (RSS).

scREAD consolidates a variety of web frameworks to provide user-friendly interactive visualizations. The front end was built on top of Nuxt.js (https://nuxtjs.org/) and utilized libraries such as Vuetify (https://vuetifyjs.com/en/) and Plotly.js (https://plotly.com/). Koa.js (https://koajs.com/) serves as the REST API back-end server for data query and custom job submission. All data are stored and managed using a MySQL database. The entire web application is containerized with Docker and deploys on a Red Hat Enterprise seven Linux system with 28-core Intel Xeon E5–2650 CPU and 64GB RAM. All integrated tools are listed in Supplementary Table S6.

## 3 Conclusion and discussion

In this paper, we describe the first release of scREAD, which is the first database collecting all existing human and mouse scRNA-Seq and snRNA-Seq datasets with AD pathology and provides a one-stop interactive visualization of the control atlas and analysis results based on these datasets. These datasets have been published and freely accessible in the public domain as of June 1st, 2020. The number of cells and the statistical distribution of these 55 datasets are shown in Supplementary Figure S1,2. Furthermore, scREAD allows users to submit a new dataset to reproduce all the analyses results showcased in scREAD in support of their AD research. We believe that our database would benefit the AD researchers through studying the data and corresponding analysis results in scREAD in particular.

In the future, we will continue to collect more AD scRNA-Seq & snRNA-Seq data from more brain regions, and build up healthy atlas in diverse brain regions of human, mouse and extend to other species. Meanwhile, we will collect AD single-cell omics data, such as scATAC-seq and proteomics data, and achieve more comprehensive analysis results based on single-cell multiple omics data. In addition, spatial transcriptomics and In situ sequencing have been recently used in studying AD (Chen, et al., 2020). The transcriptome-scale spatial gene expression datasets can further provide insights in answering the regional and cellular vulnerability in AD. Thus, we will specifically add the spatial transcriptomics and In situ sequencing datasets from human AD and AD-like animals to the current scREAD to enable more functional interpretation.

## Acknowledgements

We thank Yang Li, Zhenyu Wu, Liangping Li, Shane Chen, and Lalitha Venkataraman for their assistance in database testing.

## Funding

This work was supported by the National Institutes of Health [R01 GM131399-01, Q.M.; K01 AG056673, H.F.], the Department of Defense [W81XWH1910309, H.F.], the Alzheimer’s Association [AARF-17-505009, H.F.], and the Neuroscience Research Institute Pilot Award and the Chronic Brain Injury Pilot Award from The Ohio State University [H.F.]. The content is solely the responsibility of the authors and does not necessarily represent the official views of the National Science Foundation and the National Institutes of Health.

## Conflict of Interest

none declared.

## Notes

### Competing Interest Statement

The authors have declared no competing interest.

