## Supplementary material for "scREAD: A single-cell RNA-Seq database for Alzheimer’s Disease": scread_supplementary.docx

^3^ School of Aerospace Engineering, Xiamen University, Fujian, China,

^4^ School of Computer Science and Technology, College of Intelligence and Computing, Tianjin University, Tianjin, China

Tutorial of scREAD

Supplementary Table S1: The dataset source.

Supplementary Table S2: The brain regions are covered in scREAD for human and mouse species.

Supplementary Table S3: Information of tested data.

Supplementary Table S4: The marker genes used to assign a cell to a cell type.

Supplementary Table S5: The selection of differential gene expression analysis between different conditions (Condition 1 v.s. Condition 2) for diverse cell types in scREAD.

Supplementary Table S6: The computational tools used in scREAD.

Supplementary Figure S1: The number of cells in each of 55 files.

Supplementary Figure S2: The distribution of the species, gender, condition, and brain region for 55 files.

**Tutorial of scREAD**

The scREAD database includes six parts, and they are Home, Browse control atlas, Submit, Help, Download, and the steps to run backend workflow locally. The 'Home' page contains four statistical pie charts that reflect ratio distribution in 55 datasets for each of the four factors (species, condition, region, and gender), and a table that contains the information of these 55 datasets. The 'Browse control atlas' page includes the 15 control atlases from different brain regions of human and mouse species. The 'Submit' page provides an interface that can let users submit their AD scRNA-Seq & snRNA-Seq datasets into scREAD to do the same analysis as shown in our database. The 'Help' page contains three modules, e.g., Frequently asked questions, Usage, and Contact, and users can find the corresponding information for each of these three modules. The 'Download' page provides downloads for the datasets that are stored in scREAD. The ‘Running backend workflow locally’ part provides the scripts of the scREAD workflow.

### **Part 1. Home page**

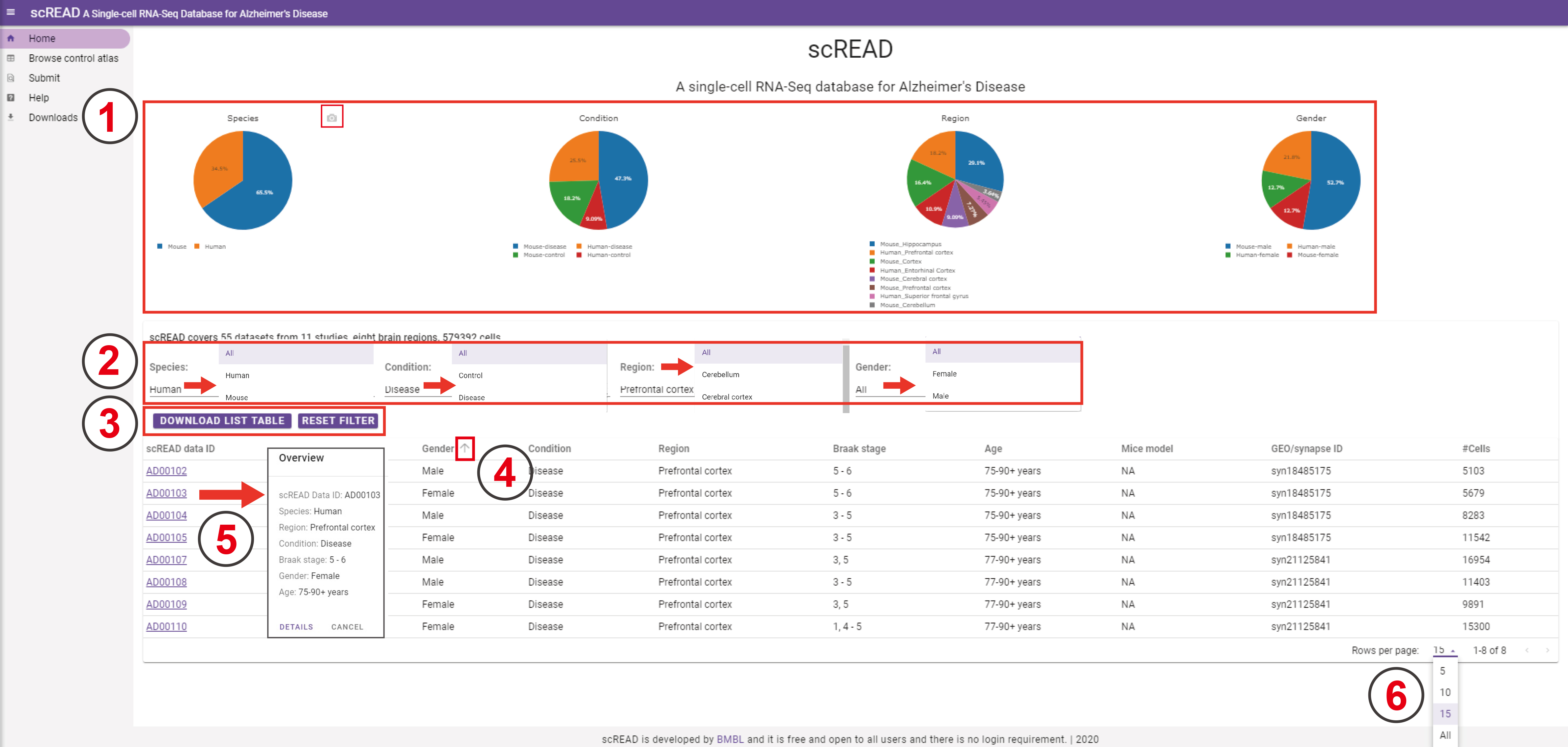

1. Here is the general statistical information of all scRNA-Seq & snRNA-Seq datasets that are covered in scREAD. The pie charts represent four factors of distribution: species, control/disease condition, brain region, and gender from the left side to the right side, respectively. Each color in each pie chart represents one element, and the number represents the distribution ratio for each element under each factor for 55 datasets. The scREAD contains a total of 579,392 cells across eight different brain regions of human and mouse species. The pie charts can be downloaded by clicking the camera icon on the top right of each pie chart.

- For the pie chart of the species, there are 19 human species datasets and 36 mouse species datasets across 55 datasets.
- For the pie chart of the control/disease condition, there are 5, 14, 10, and 26 datasets across 55 datasets corresponding to human control, human disease, mouse control, and mouse disease, respectively.
- For the pie chart of the brain region, there are 6, 10, 3, 9, 2, 5, 16, and 4 datasets across 55 datasets corresponding to human entorhinal cortex, human prefrontal cortex, human superior frontal gyrus, mouse cortex, mouse cerebellum, mouse cerebral cortex, mouse hippocampus, and mouse prefrontal cortex regions, respectively.
- For the pie chart of the gender, there are 12, 7, 29, and 7 datasets across 55 datasets corresponding to for human male, human female, mouse male, and mouse female, respectively.

2. Here are all the options for the species, control/disease condition, brain region, and gender factors. The following table is the whole table of the general information across 55 datasets. The default display is the whole table, and the human dataset's information appears on the top. It will show the searching result table based on users' choices. Users can select the 'All' option, and it will return search results containing all elements. Users can select an option, and it will return search results including only entries matching the option selected.

- Note: When users choose one of these factors, the corresponding search results will be changed based on the factors that the users make. Users can select different factors simultaneously, and the return result is the intersection of all the options that they choose.

3. Users can download the current table, which is based on the options that users made, by clicking the 'DOWNLOAD LIST TABLE' button, and they can return to the default show by clicking the 'RESET FILTER' button.
4. For each of these nine factors (Species, Gender, Condition, Region, Braak stage, Age, Mice model, GEO/synapse ID, and #Cells), users can achieve the table that is sorted on increasing/decreasing order, by clicking this button or just click the name of factors. Note: This button will show up when the cursor is moved on top of the name of these factors.
5. When users click the scREAD data ID, a floating window will show up. In this floating window, there is an overview of information for this dataset. Users can click the "DETAILS" button and scREAD will go to the analysis result page to browse the analysis results of this dataset. Users can also click the "CANCEL" button to exit and choose another dataset to browse. If users don't click the 'CANCEL' button, it will automatically go to the analysis result page in ~ 5 seconds.
6. This is how many rows of the whole table will show on the home page. The default number is 15, but users can choose another number in the options bar.

### **Part 2. Example result illustration**

We used the dataset of [AD00103](https://bmbls.bmi.osumc.edu/scread/AD00103) as an example to show the analysis result. This dataset is consisting of 6,629 cells isolated from human AD female prefrontal cortex ([Mathys et al., 2019](https://pubmed.ncbi.nlm.nih.gov/31042697/)).

This tutorial will guide you through the analysis result page of scREAD in detail.

### 2.1 General information

The general information of this dataset is shown in the first section of the analysis result page. This section includes four parts: Overview, Dataset information, Dataset source, and Datasets from the same experiment.

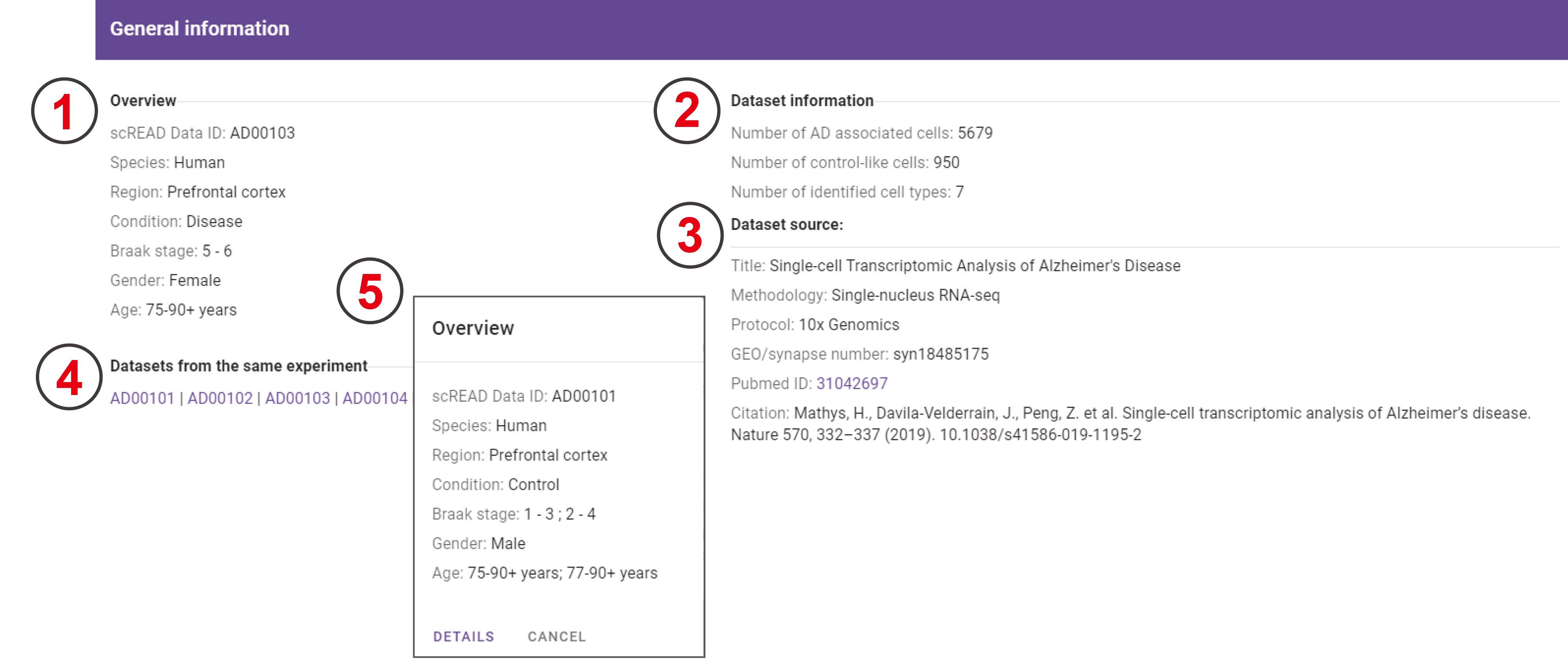

1. The "Overview" part is the overall detail information of this dataset. It includes seven fields of the current dataset: 'scREAD Data ID', 'Species', 'Region', 'Condition', 'Braak Stage', 'Gender', and 'Age'. Here's the overview information of 'AD00103' dataset.
2. The "Dataset information" part contains three entries: the number of cell types that were identified in this dataset, how many control-like cells, and how many AD-associated cells.
3. The "Dataset source" part is the general information of the corresponding research paper for this dataset. It includes the title of this paper, the scRNA-Seq/snRNA-Seq technique used in this paper, protocol, GEO/syn ID, Pubmed_ID, and how to cite this paper.
4. In the "Datasets from the same experiment" part, it lists all the datasets that are included in the same experiment. All the datasets shown here come from the same paper, and they usually contain the control and disease datasets.
5. When users click the name of the scREAD Data ID, a window that includes the overview of this dataset to select will appear. Then users can click the 'DETAILS' button to go to the analysis result page of the corresponding dataset. Users can also click the 'CANCEL' button, and then the floating window will disappear.

### 2.2 Cell clustering

Two high-resolution UMAP plots show the predicted cell types and the expression distribution of all genes in this dataset, respectively.

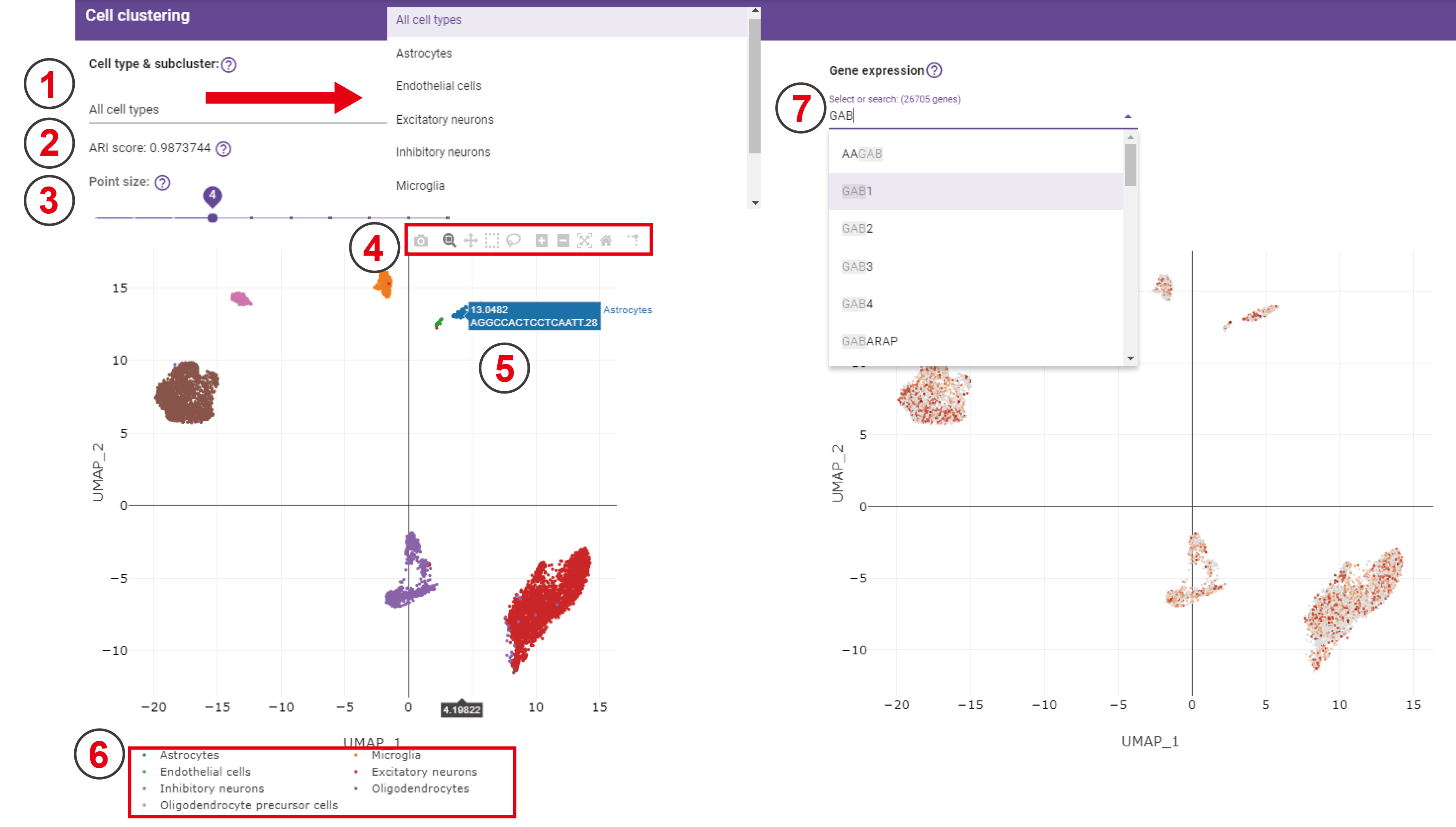

1. There are seven predicted cell types, which are shown in the drop-down bar. If users choose one of these cell types, the following UMAP will change to the UMAP of predicted subclusters for this specific cell type.
2. The ARI score is used to evaluate the performance of our predicted cell types compared with the original cell labels from the original paper. The higher the score is, the more consistency there is between our predicted cell labels and the original cell labels. Note: If we don't have the ARI score, it will show a silhouette score instead.
3. A sliding bar is used for controlling the size of each point in the following UMAP. It ranges from 1 to 10, the bigger the number is, the larger the point size is.
4. This function bar contains several quick buttons for graphic operations. For example, the first button is 'Download plot as a png', users can click this button and this UMAP figure will be downloaded to the user’s local computer automatically. For the 'Zoom in' and 'Zoom out' buttons, users can zoom in or zoom out this figure by clicking these two buttons. If users want to return to the default figure after they applied zoom in or zoom out in this figure, users can click the 'Reset axes' button to let this figure go back to the default status.
5. A floating window will show up when users move their cursor on cells, indicating the belonging cell type, cell name, and the UMAP coordinate.
6. This is the legend of this UMAP.
7. The genes in the drop-down bar are all genes expressed in this dataset, and users can also input the name of genes that they're interested in. The darker the color is in this UMAP, the higher the expression value of the gene. The function bar of this UMAP is the same as the cell type UMAP.

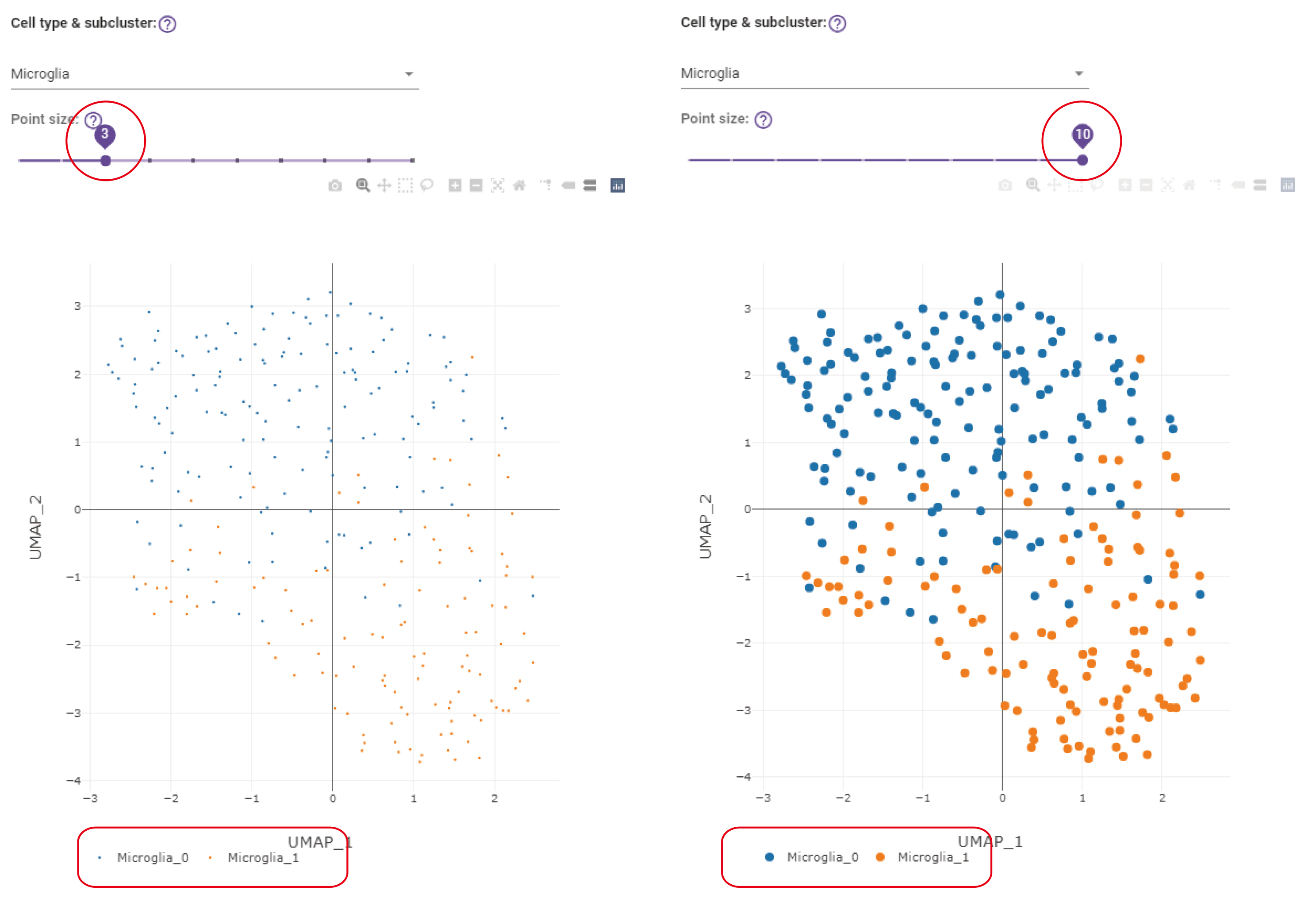

It is a UMAP of the predicted subclusters for microglia cell type. There are two predicted subclusters for microglia, e.g., microglia_0 and microglia_1. The point size of this UMAP can be adjusted by sliding left and right through the sliding bar. The default point size is 4. For the UMAP at the left panel, the point size set as 3. For the UMAP at the right panel, the point size is set as 10. The UMAP of subclusters is clearer after increasing point size.

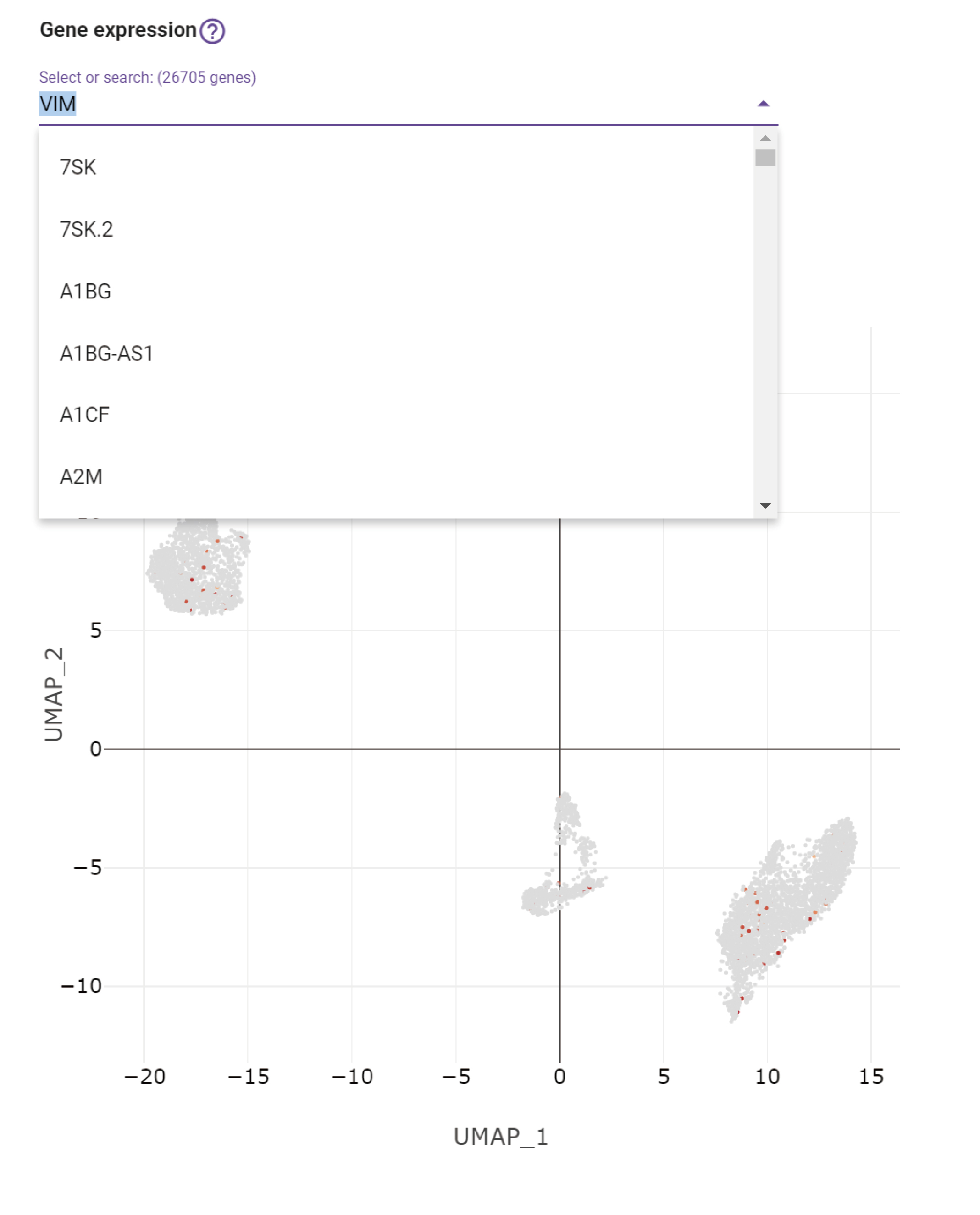
 The genes in the drop-down bar are all of the genes expressed in this dataset and ranked in alphabetical order.

### 2.3 Differential expression (DE) / Gene set enrichment

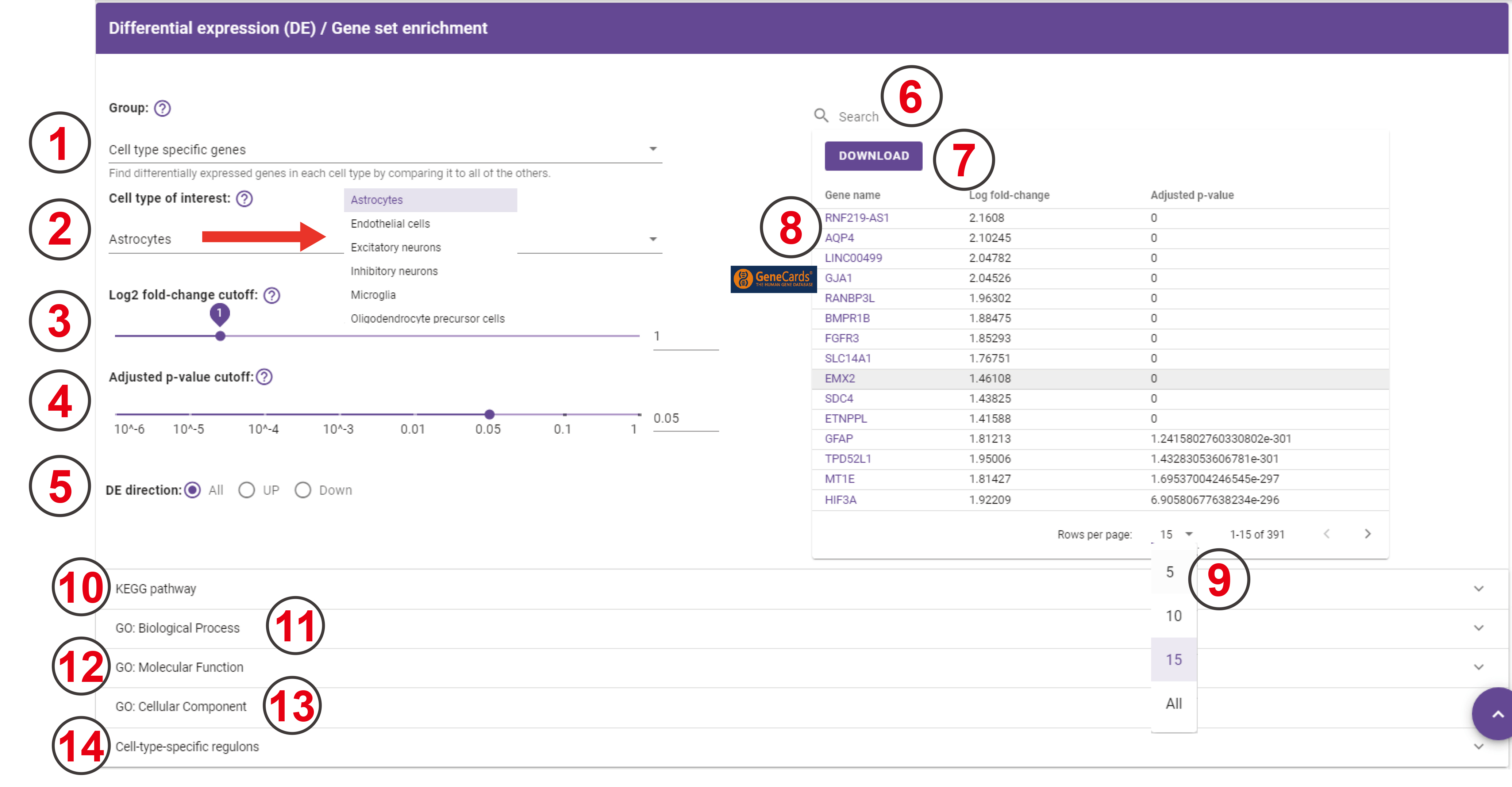

1. There are five groups here. The default is 'Cell type specific group'. And the following choice will be changed as users choose different groups. For the current dataset, users can achieve the cell-type-specific differentially expressed genes for each predicted cell type, and can also achieve the specific differentially expressed genes for each predicted subcluster. Users can also achieve the cell-type-specific differentially expressed genes across different datasets.
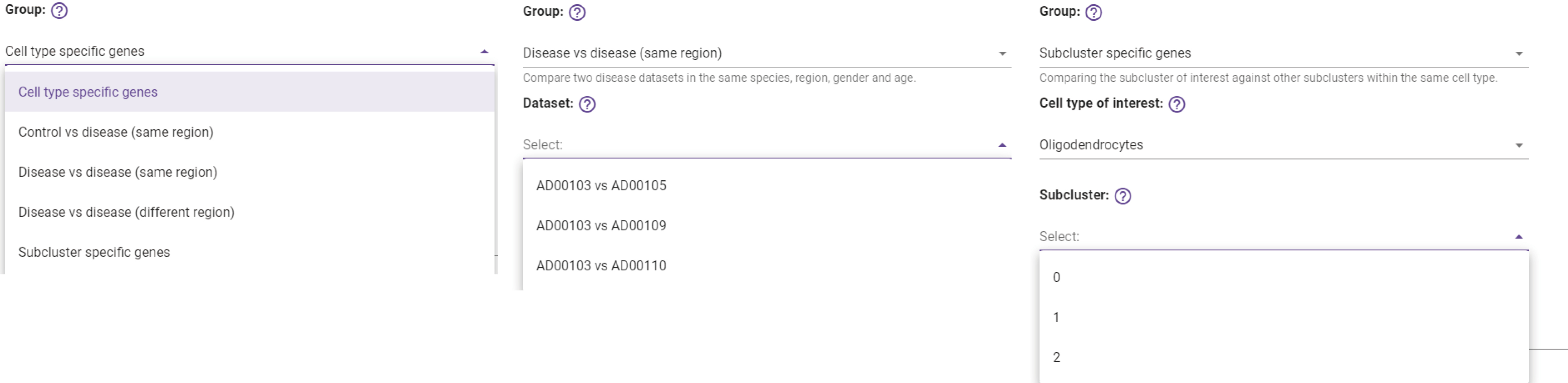

2. Here users can choose all the cell types that are predicted with the "Cell Clustering" section. There are seven predicted cell types in the drop-down bar, users can choose one cell type to achieve the cell-type-specific differentially expressed genes.
3. The Log2 fold-change ranges from 0 to 5.
4. The Adjusted p-value ranges from 10^-6 to 1.
5. The DE detection contains three options, all, up, and down.
6. Users can search for genes that they are interested in, and then the following table will return the matching result.
7. Users can use this download button to download the current table.
8. [GeneCards](https://www.genecards.org) database is linked to each gene in the table.
9.
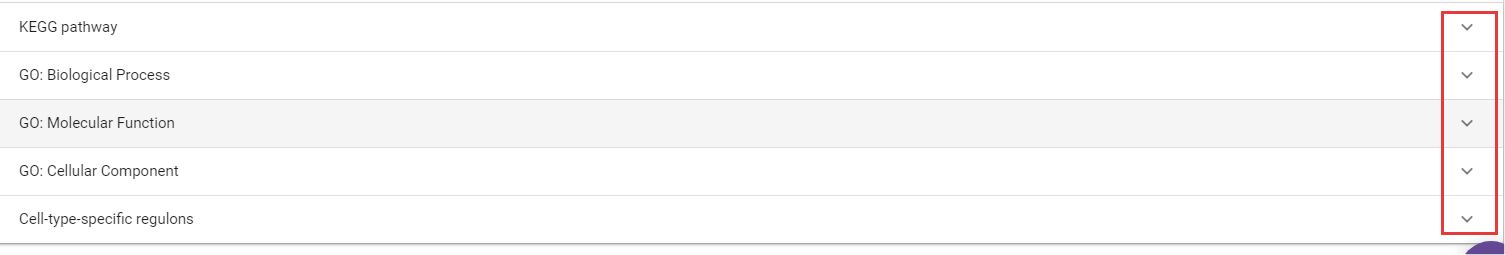
This is how many rows of the table that will show. The default number is 15, but users can choose another number in the options bar, e.g. 5, 10, 15, and all.
10. KEGG pathway enrichment analysis result table of the DEGs will appear when users click the inverted triangle, and this table can be downloaded when they click the 'Download' button. When users click the reversed triangle at the end of each row in this table, it shows the genes that are enriched on this pathway, and this table can be downloaded when they click the 'Download' button. Users can also search for a specific item by entering the content they want to search in the search box.
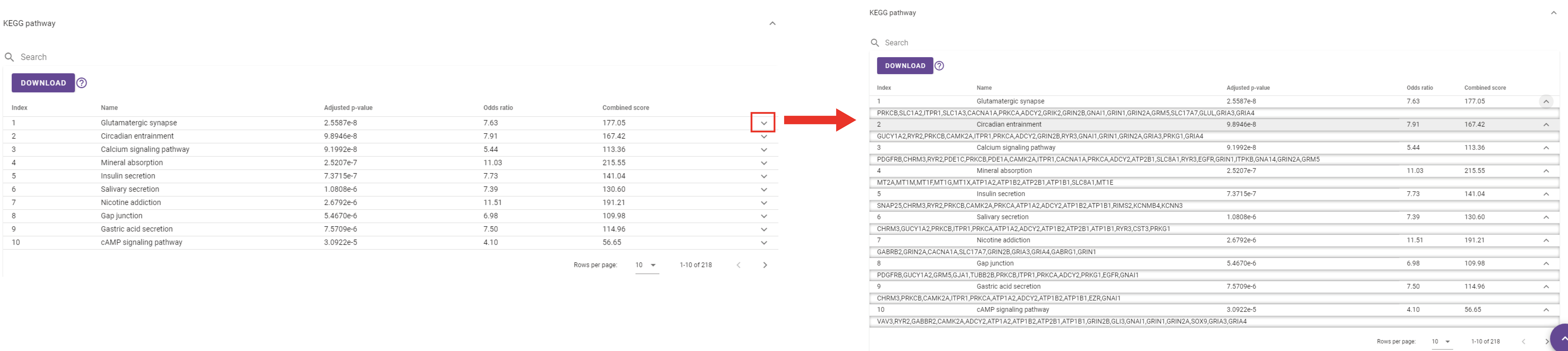

11. GO biological process analysis result table of the DEGs will appear when users click the inverted triangle, and this table can be downloaded when they click the 'Download' button. When users click the reversed triangle at the end of each row in this table, it shows the genes that are enriched on this item, and this table can be downloaded when they click the 'Download' button. Users can also search for a specific item by entering the content they want to search in the search box.
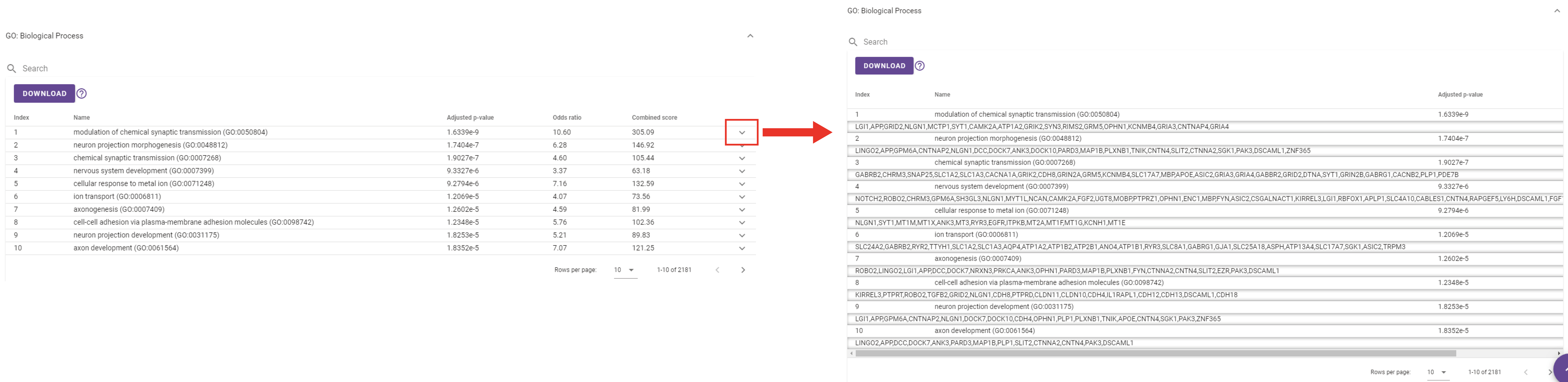

12. GO molecular function analysis result table of the DEGs will appear when users click the inverted triangle, and this table can be downloaded when they click the 'Download' button. When users click the reversed triangle at the end of each row in this table, it shows the genes that are enriched on this item, and this table can be downloaded when they click the 'Download' button. User can also search for a specific item by entering the content they want to search in the search box.
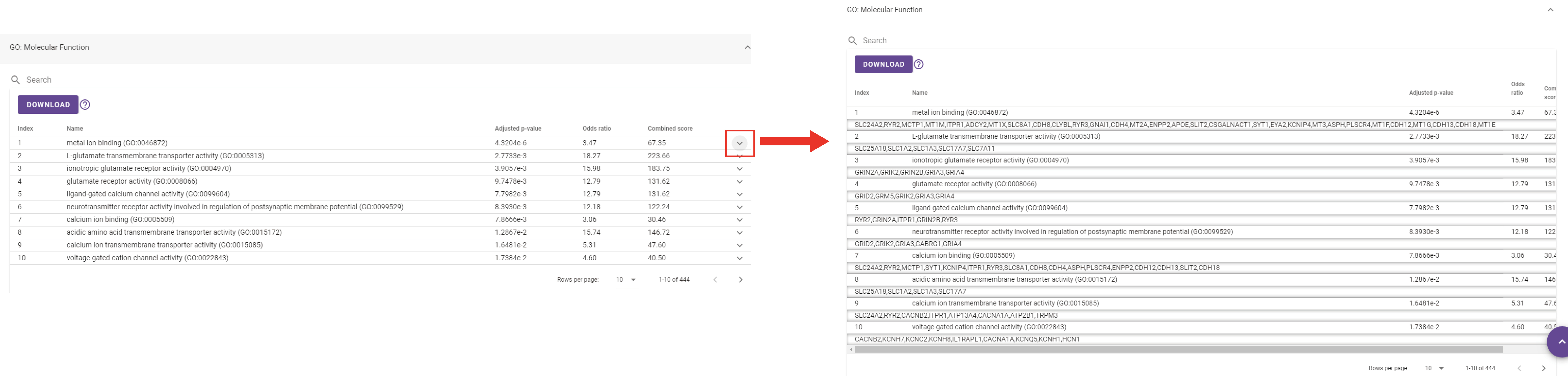

13. GO cellular component analysis result table of the DEGs will appear when users click the inverted triangle, and this table can be downloaded when they click the 'Download' button. When users click the reversed triangle at the end of each row in this table, it shows the genes that are enriched on this item, and this table can be downloaded when they click the 'Download' button. Users can also search for a specific item by entering the content they want to search in the search box.
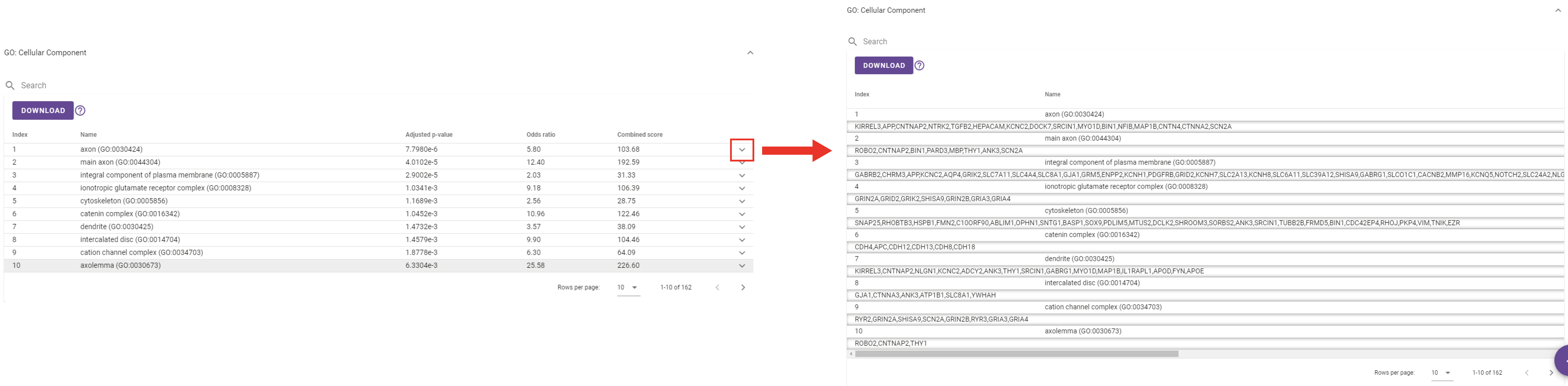

14. Cell-type-specific regulon analysis result table of this dataset will appear when users click the inverted triangle, and this result only shows up when they choose the 'Cell type specific genes' in the 'Group' drop-down bar.

-
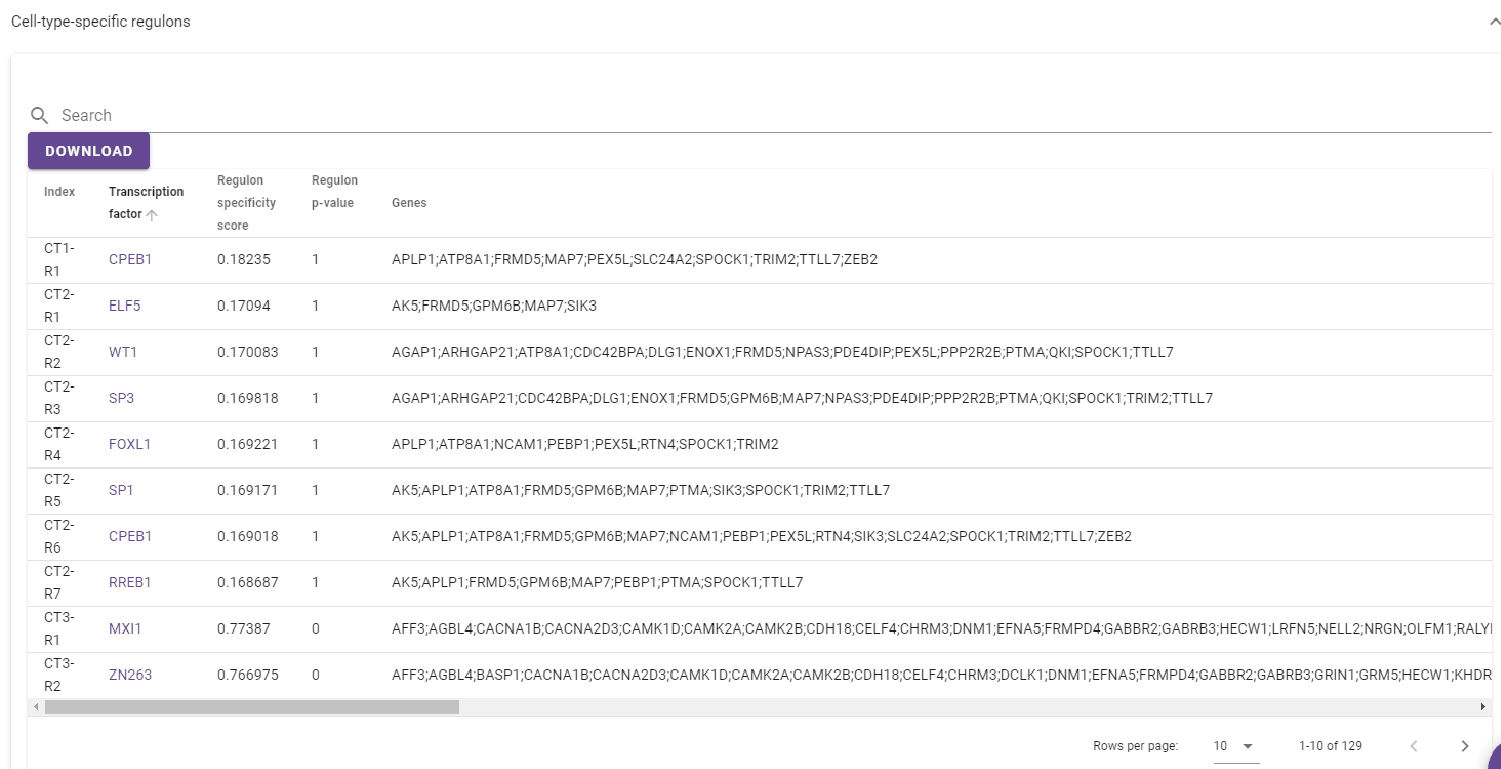
 This is the cell-type-specific regulon result table for each cell type, and this table can be downloaded when users click the 'Download' button.
- Regulon heatmap
-
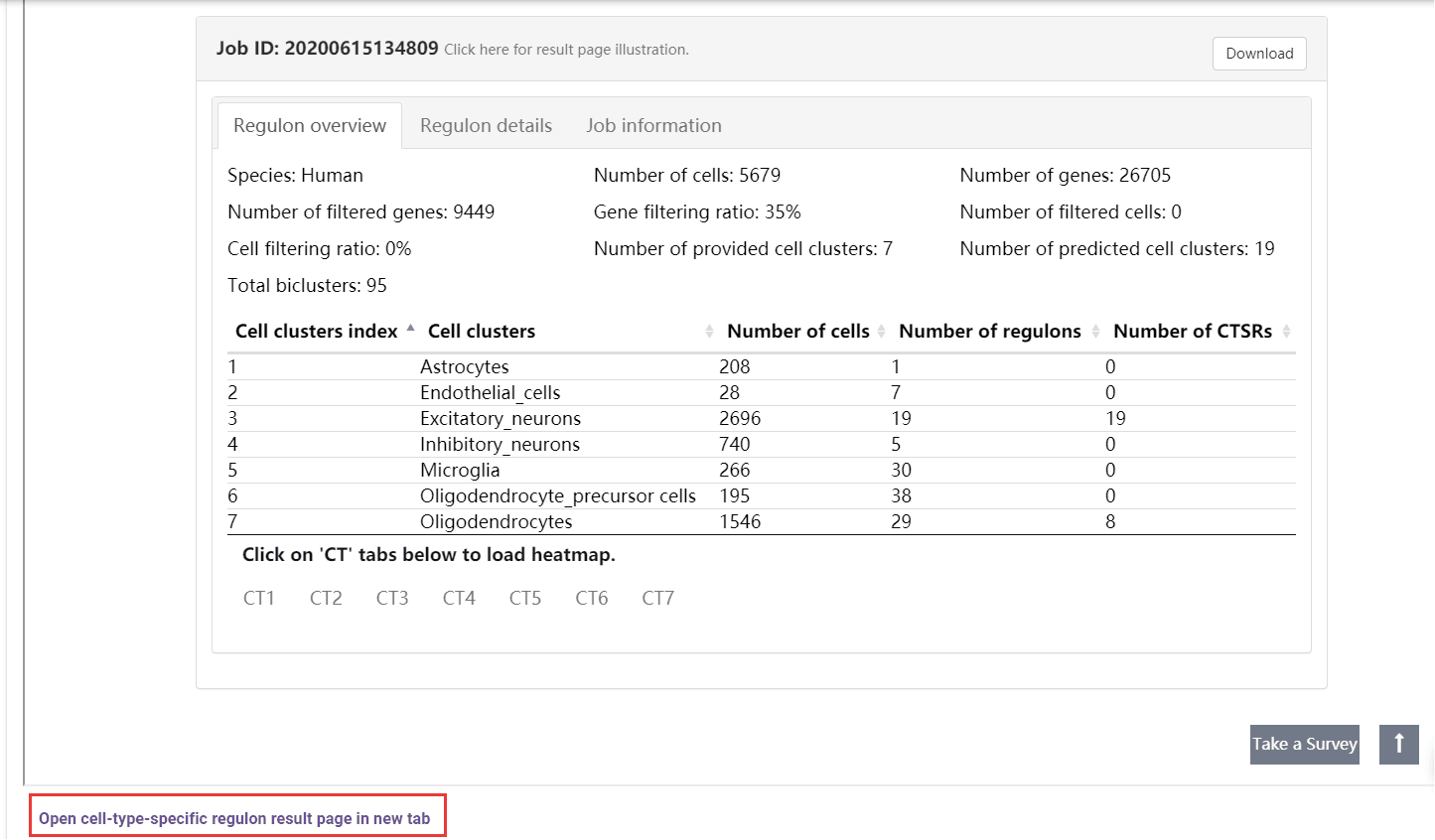
 A table summarizes the clusters shows the overall cell number and regulon number in each cluster. Users can jump into the [IRIS3](https://bmbl.bmi.osumc.edu/iris3/results.php?jobid=20200615134809) to see the detailed results of this job by clicking the 'Open cell-type-specific regulon result page in the new tab' button. In the table, the index number will be given to represent CTSRs. A cell-gene-regulon heatmap will be displayed for each cell type by clicking on the CT# tab. See the figure details below.
-
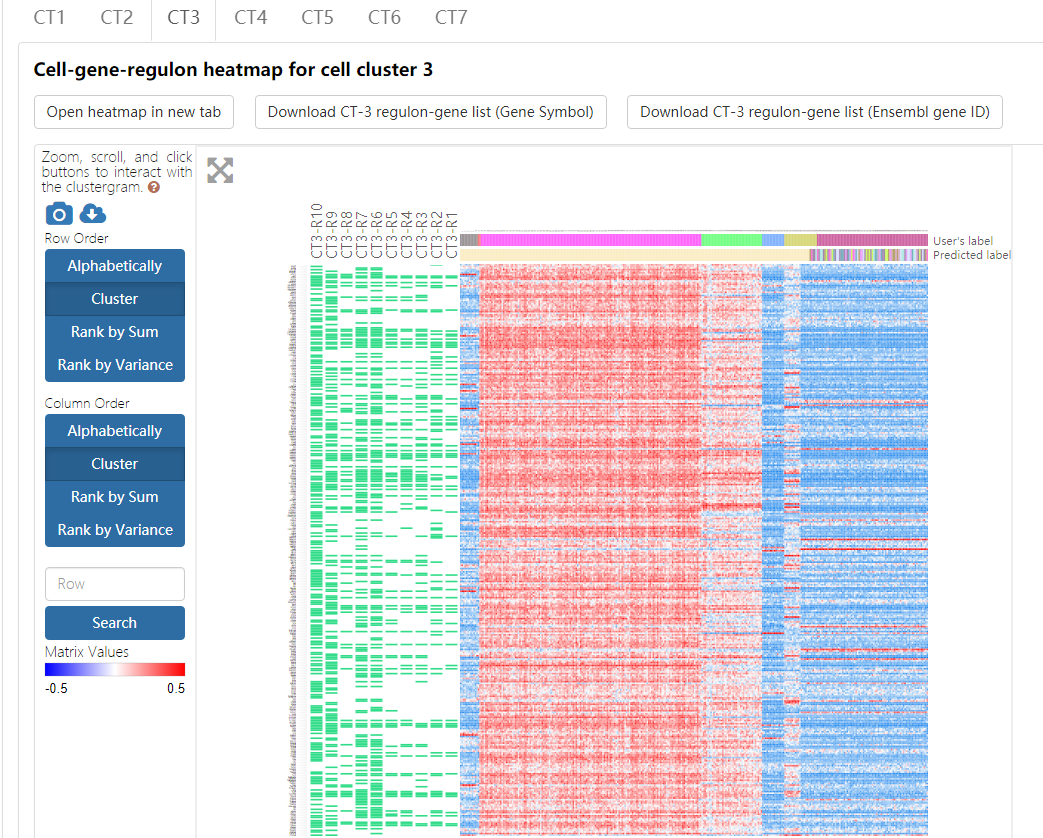

- The heatmap, empowered by Clustergrammer, showcases the expression pattern of genes from the top ten regulons in the corresponding cell type. To explore the details about Clustergrammer, please read the [Clustergrammer documentation](https://clustergrammer.readthedocs.io).
  - Both gene compositions of regulons and their expression values across different cell types can be intuitively displayed in such a heatmap. Regulons are ranked in increasing order of the empirical p-values of regulon specificity scores (RSS) as described above, and a regulon is named as CTn-Rm with n representing the index of cell type and m represents the regulon rank. Due to the space limitation, only the top ten regulons and their corresponding genes are showcased in the heatmap, and the component genes of each regulon are indicated as green rectangles. The heatmap records the log-transformed expression level of each top-ten-regulon-covered gene across all cells.
  - Cell names, original cell type labels, and cell types labels predicted by Seurat are shown on the heatmap. This heatmap can also be sorted by gene and cell by double-clicking on the appropriate area on the image. Conveniently, a series of gene enrichment tests can be directly performed on the heatmap using the integrated Enrichr function in the Clustergrammer framework.
  - This heatmap can be downloaded by clicking on the top-left buttons above the heatmap and choosing the 'Take a snapshot' option.
- Regulon details
-
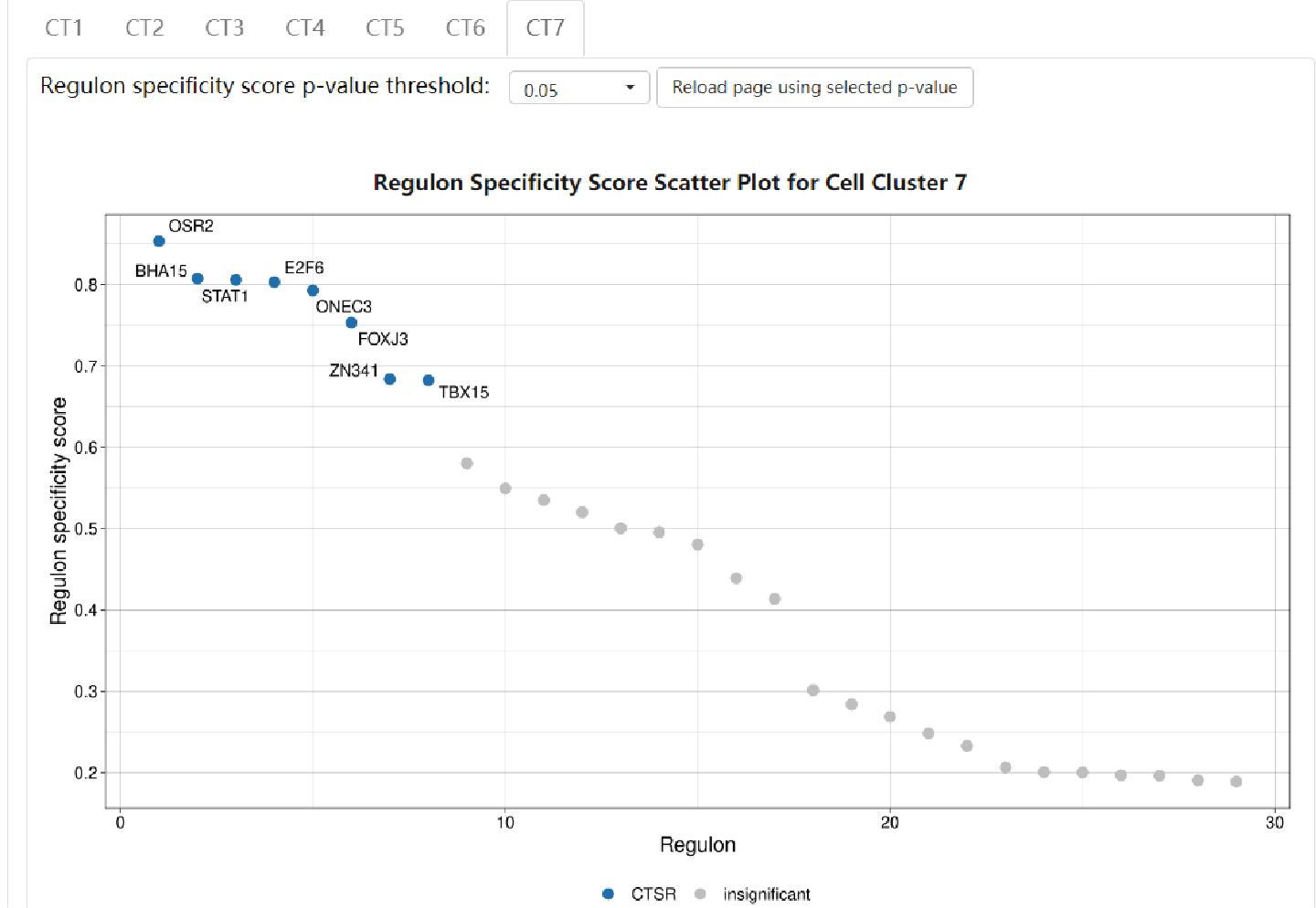

Regulon results are separately showcased in each cell type. Click on the "CT#" button to switch to see results in other cell types. A scatter plot shows the distribution of RSS of each regulon. CTSRs are marked as blue dots with their representative TF names beside, and insignificant regulons are marked as grey dots. Detailed information of each regulon can be found in [IRIS3](https://bmbl.bmi.osumc.edu/iris3/index.php).

### **Part 3. Browse control atlas page**

The 'Browse control atlas' page contains all the 15 control atlases that are stored in the scREAD based on different brain regions for different species and different mouse ages.

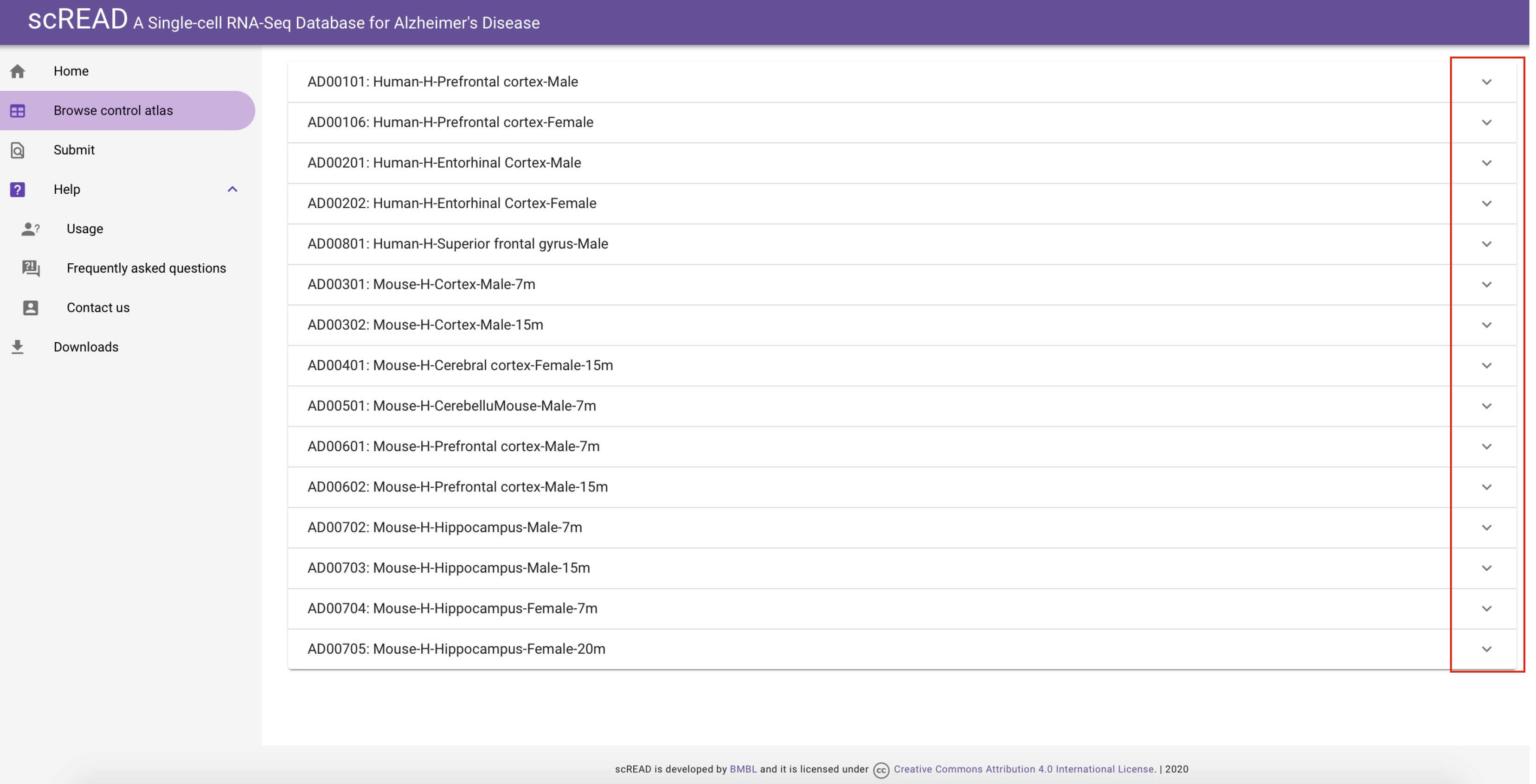

These are 15 control atlases entries. The default pattern is all the UMAP of control atlases are folded, however, users can click the reverses triangle to unfold the UMAP of each control atlas. The top 5 control atlases are human control atlases, and the rest control atlases are mouse control atlases.

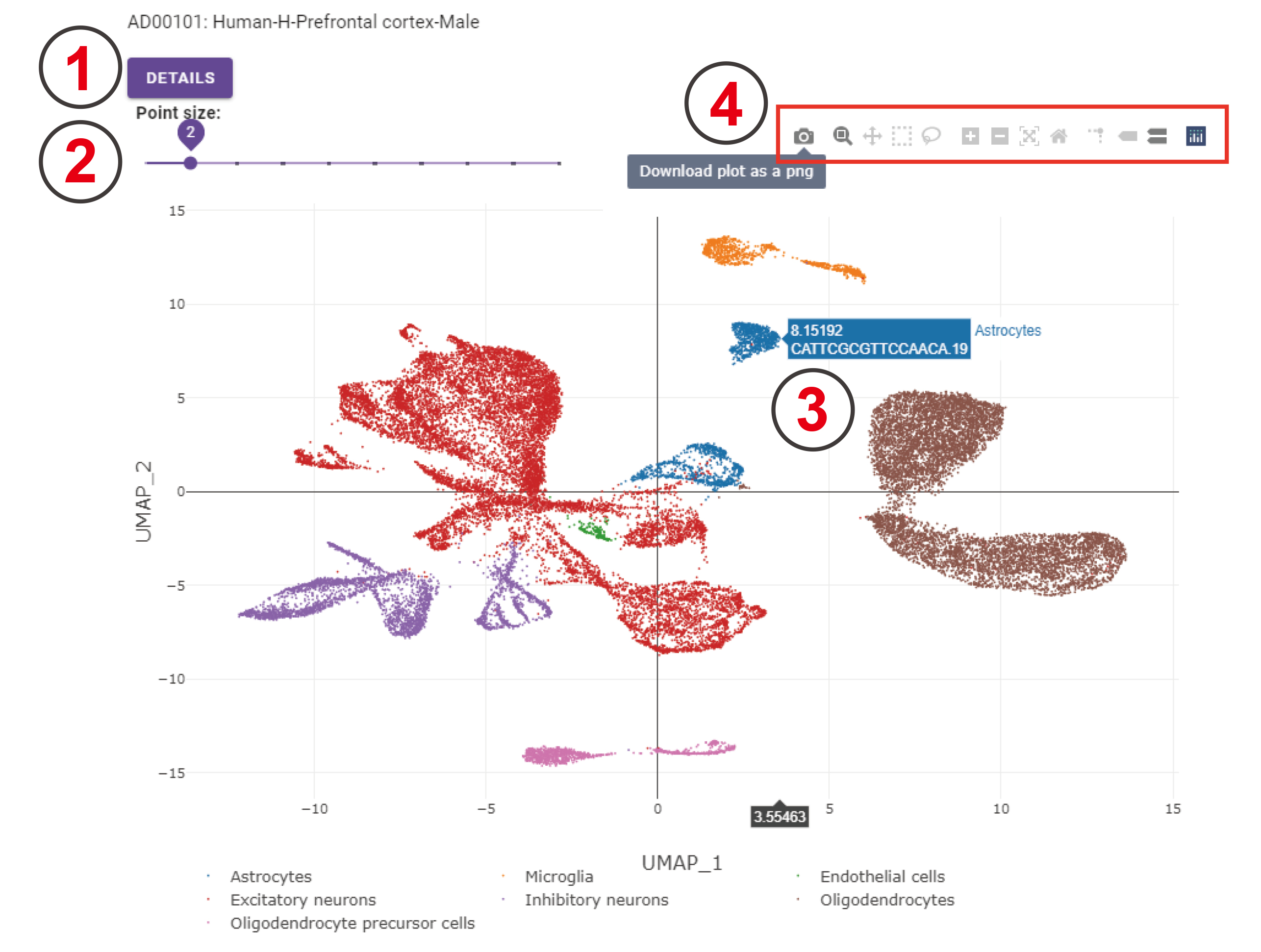

This is the UMAP of control cells coming from the prefrontal cortex region of human male datasets. 1. Clicking the 'DETAILS' button will navigate to the analysis result page of this dataset. 2. The point size on this UMAP can be adjusted by clicking the point on this sliding bar. The point size ranges from 1 to 10. 3. A floating window will appear when moving the cursor on cells, indicating the belonging cell type, cell name, and the UMAP coordinate. 4. This bar has 11 options; users can choose each option by clicking it and the corresponding UMAP will be changed as users change the options. For example, users can download this UMAP by clicking the 'Download plot as a png' button.

### **Part 4. Submit page**

The submission of new entry is welcome, and it can be done on the “submit” page. One scRNA-Seq file of AD disease should be uploaded, and one scRNA-Seq file of control can be uploaded or not.

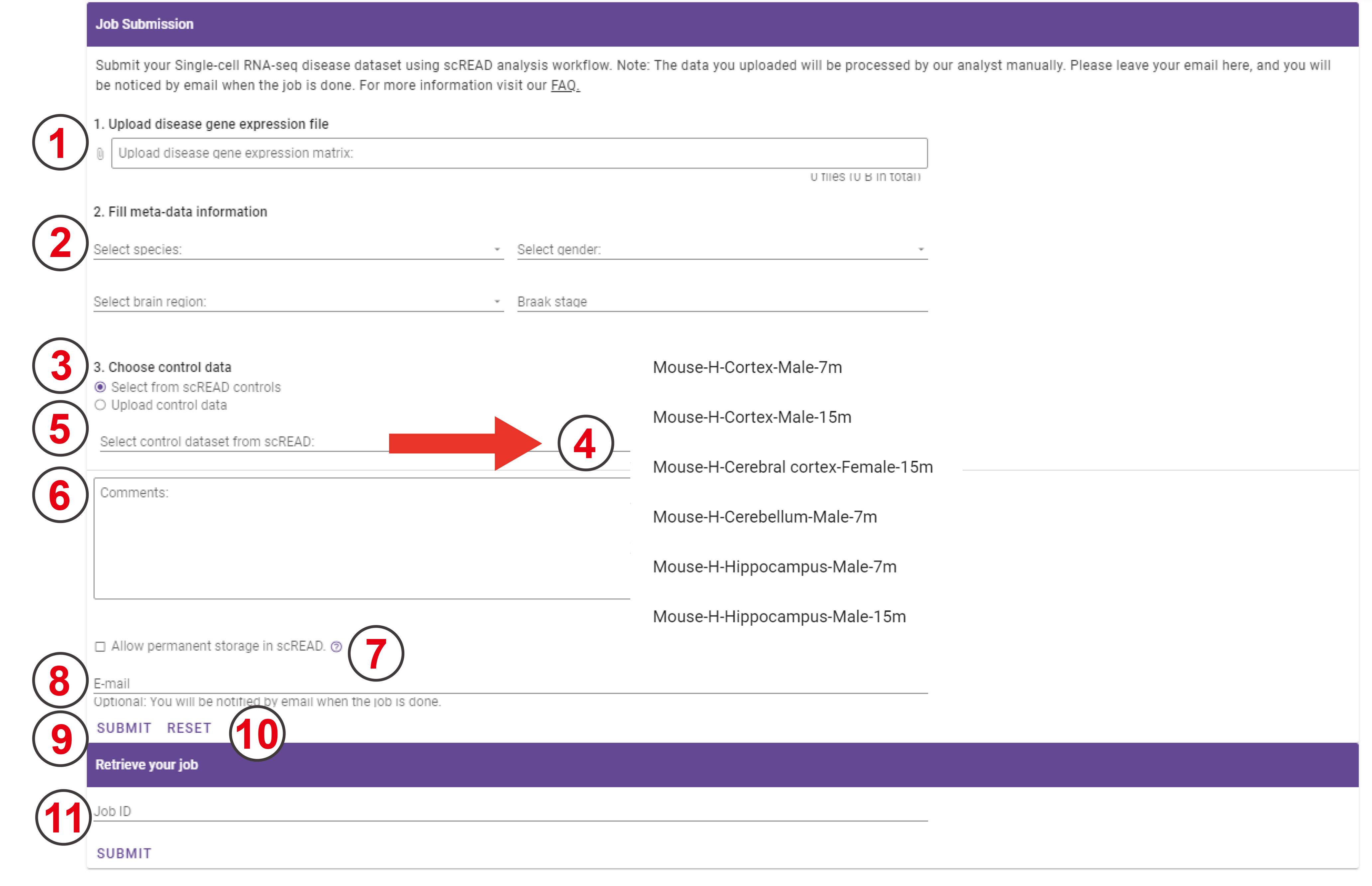

1. Upload your AD scRNA-Seq expression matrix file by selecting the file stored on your computer. Note: This file is required if you want to analyze new data. Note: The format of your uploaded file should be a text format.
2. You can provide species, gender, brain region, and Braak stage these four types of information of your input gene expression dataset to scREAD.
3. You can select one of the control datasets as a reference control atlas to do the downstream analysis by choosing 'Select from scREAD controls' option.
4. These are all the 15 control files that are stored in scREAD to produce all control atlas. For human control files, the first column is the species, the second column is the condition, the third column is the brain region, the last column is the gender. For mouse control files, the first column is the species, the second column is the condition, the third column is the brain region, the fourth column is the gender, and the last column is the age of the mouse.
5. You can also upload your control dataset if you have one to do the comparison within your own paired dataset by selecting 'Upload control data' and then click the bar.
6. If you have any comments about scREAD, we will be appreciative that you can write your comments here.
7. Clicking this option, it means you allow us to store your data in scREAD (both datasets and results) for the future database construction. Be cautious if your data have not been published.
8. An email is not required to submit the job; however, we strongly suggest you provide your email because the data you uploaded will be processed by our analyst manually. So you will be noticed by email when the job is done.
9. Submit the job once everything is ready. If you have provided your email to us you will receive an email after you submit your job successfully. The job ID is in the showed up floating window, which can be used to retrieve the results.

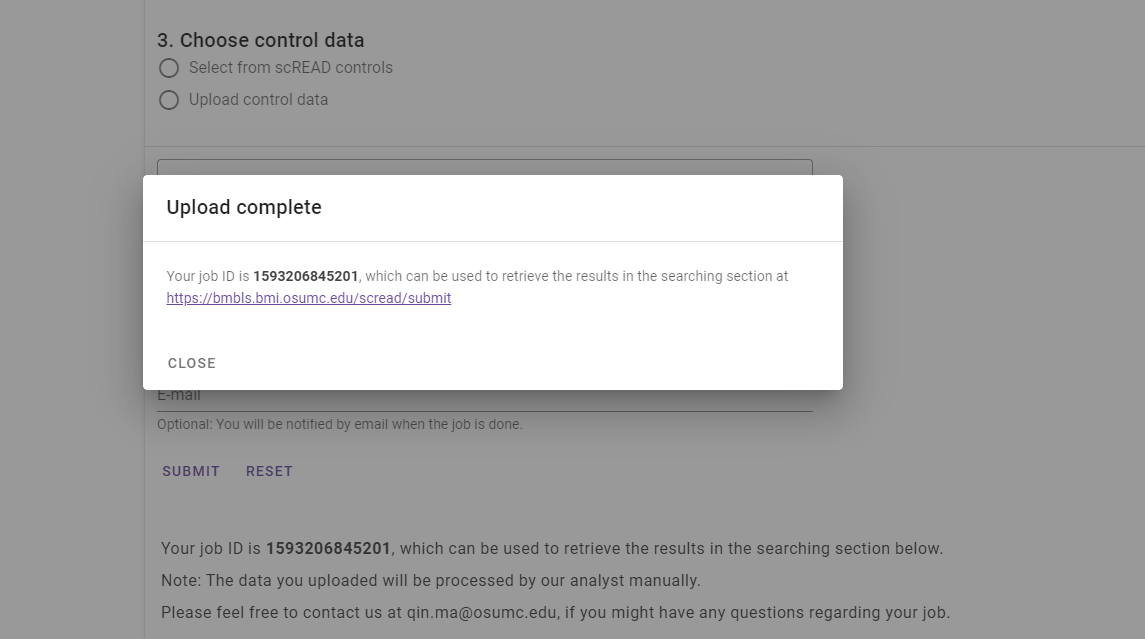

10. You can reset all the input information by clicking this button and restart over again.
11. You can input the job ID here and retrieve the analysis result after the work is done.

### **Part 5. Download page**

Not all the datasets in scREAD are available to download for users. On the “Download” page, datasets that downloaded from the GEO database are available to download, but datasets downloaded from Synapse are not available to download.

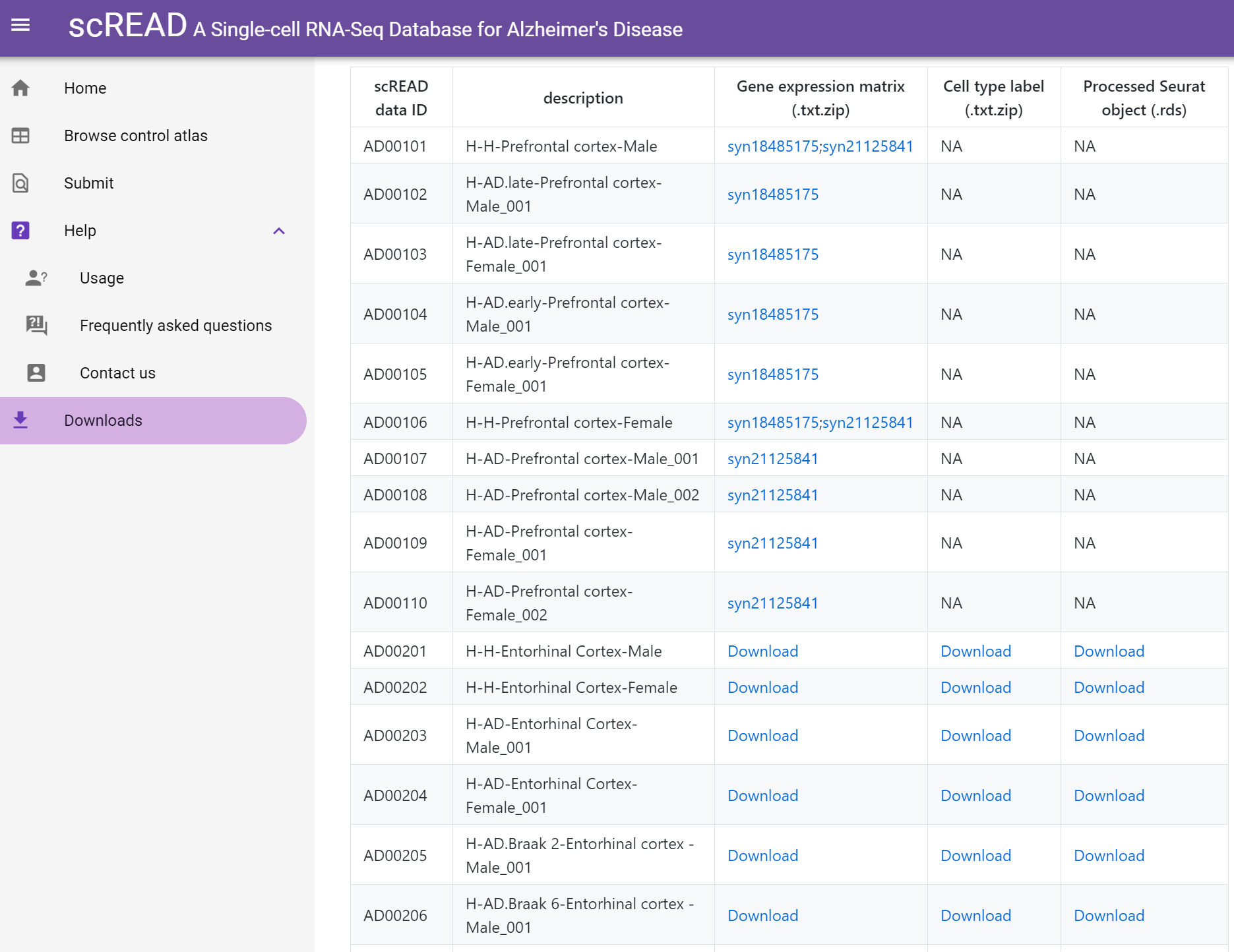

It provides three files for users to download. 1. The compressed gene expression matrix (.txt.zip); 2. Cell type labels (.txt.zip); 3. Processed Seurat R object (.rds).

### **Part 6. Running backend workflow locally**

scREAD is an open-source initiative with scripts and workflows available at <https://github.com/OSU-BMBL/scread-backend> and the frontend code repository available at <https://github.com/OSU-BMBL/scread>.

The workflow in R can be found in <https://github.com/OSU-BMBL/scread/tree/master/script>, the folder contains the following files:

1. custom_marker.csv. A manually created marker gene list file used for identified cell types.
2. functions.R. Visualization functions used in R.
3. build_control_atlas.R: build control cells atlas Seurat object from count matrix file.
4. transfer_cell_type.R: filter out control-like cells in disease dataset
5. run_analysis.R: run analysis workflow, and export tables in scREAD database format.
6. example_control.fst. The example control dataset.
7. example_disease.fst. The example disease dataset.

**Build control atlas**

1. Goal: Build the control atlas file from the raw gene expression matrix.
2. Prepare your control gene expression data in fst format (<https://www.fstpackage.org/>), we used fst package to store raw data in scREAD since it provides a fast, easy, and flexible way to serialize data frames. In the data frame, the first column should be gene symbols, and other columns as cell labels. Put all code and data in a working directory. (e.g PATH_TO_WD), in this tutorial, we will run example_control.fst.
3. build_control_atlas.R takes three parameters: 1. Working directory path; 2. Control data path. 3. Output data ID
4. cd PATH_TO_WD
5. Rscript build_control_atlas.R PATH_TO_WD example_control.fst control_example
6. The output should contain four files:
   a) control_example.rds. The Seurat R object storing example control data.
   b) control_example_expr.txt. Filtered gene expression matrix.
   c) control_example_cell_label.txt. The first column is the cell name, the second column is the cell type information.
   d) control_example_umap.png. UMAP plot of example control data colored by cell types.

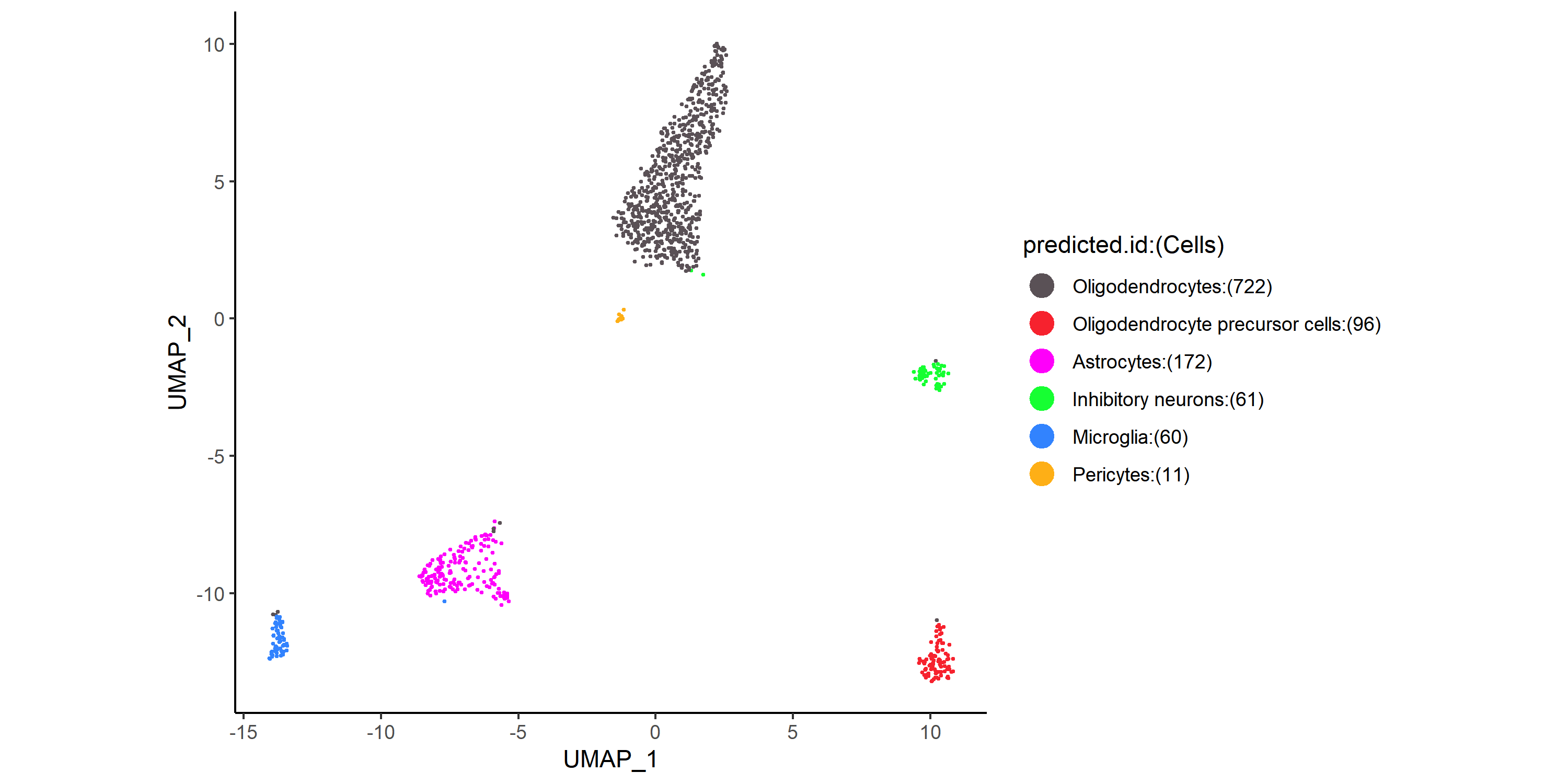

**Transfer cell types based on control atlas**

1. Goal: Annotate cell type using control atlas as the reference, onto the disease gene expression matrix file.
2. Put all code and data in a working directory. (e.g PATH_TO_WD), after you have generated the control atlas file (control_example.rds).
3. build_control_atlas.R takes four parameters: 1. Working directory path; 2. Control atlas Seurat object file name; 3. Disease gene expression matrix name; 4. Output disease data ID.
7. cd PATH_TO_WD
8. Rscript transfer_cell_type.R PATH_TO_WD control_example.rds example_disease.fst disease_example
9. The output should contain four files:
   e) disease_example.rds. The Seurat R object storing example disease data.
   f) disease_example_expr.txt. Filtered gene expression matrix.
   g) disease_example_cell_label.txt. The first column is the cell name, the second column is the cell type information.
   h) disease_example_umap.png. UMAP plot for both control and disease data colored by cell types.

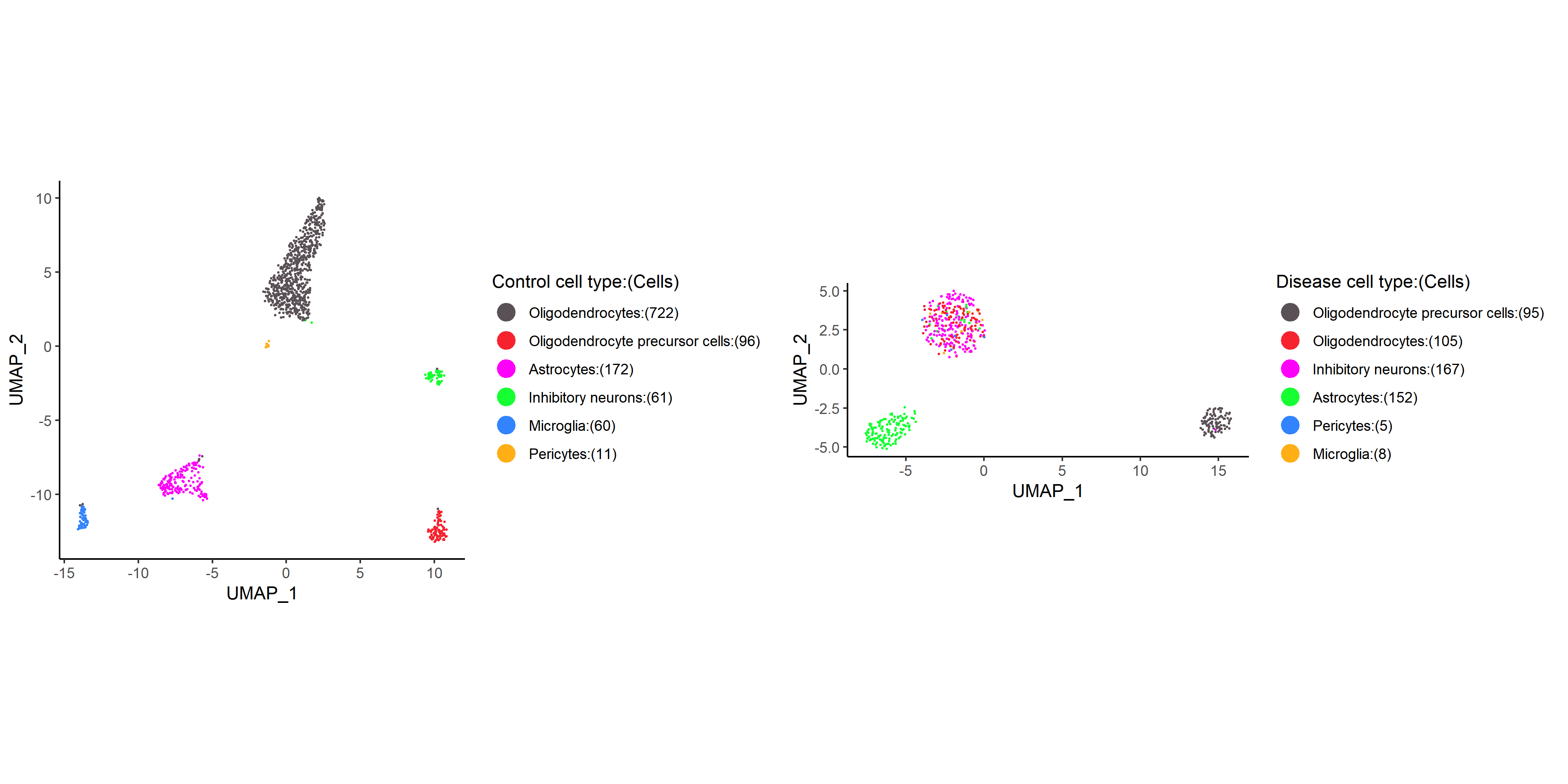

**Run data analysis**

1. Goal: Perform analysis between disease and control data
2. Put all code and data in a working directory. (e.g PATH_TO_WD), after you have generated the control atlas file (control_example.rds), and the disease file (disease_example.rds)
3. run_analysis.R takes three parameters: 1. Working directory path; 2. Control Seurat object file name. 3. Disease Seurat object file name.
4. cd PATH_TO_WD
5. Rscript run_analysis.R PATH_TO_WD control_example disease_example
6. The output should be stored in three folders:
   a) /de. Differential gene expression analysis results. 1. Cell-type-specific genes; 2. Sub-cluster specific genes; 3. Cell type DE genes between two conditions.
   b) /dimension. UMAP coordinates for two datasets.
   c) /subcluster_dimension. UMAP coordinates for each sub-clusters in two datasets.

**Supplementary Table S1**. The dataset source.

| **Species** | **Data_ID** | **Pubmed_ID** |
| --- | --- | --- |
| Human | GSE138852 | 31768052 |
| Human | syn18485175 | 31042697 |
| Human | GSE147528 | <https://www.biorxiv.org/content/10.1101/2020.04.04.025825v2> |
| Human | syn21125841 | 31932797 |
| Mouse | GSE98969 | 28602351 |
| Mouse | GSE103334 | 29020624 |
| Mouse | GSE130626 | 31902528 |
| Mouse | GSE141044 | 31928331 |
| Mouse | GSE140510 | 31932797 |
| Mouse | GSE140399 | 31932797 |
| Mouse | GSE143758 | 32341542 |
| Mouse | GSE147495 | 32320664 |

**Supplementary Table S2**. The brain regions are covered in scREAD for human and mouse species.

| **Species** | **Region** | **Brodmann area** |
| --- | --- | --- |
| Human | Entorhinal cortex | NA; NA |
| Human | Prefrontal cortex | Area 9, Area 46; Area 10 |
| Human | Superior frontal gyrus | Area 8 |
| Mouse | Cortex | NA |
| Mouse | Cerebellum | NA |
| Mouse | Cerebral cortex | NA |
| Mouse | Hippocampus | NA |
| Mouse | Prefrontal cortex | NA |

**Supplementary Table S3**. The definition of different mouse age stages in scREAD.

| **Age_Stage** | **Range of ages** |
| --- | --- |
| 2 months | 1-2 months |
| 7 months | 4-7 months |
| 15 months | 10-15 months |

**Supplementary Table S4**. The marker genes used to assign a cell to a specific cell type.

| **Cell type** | **Genes** |
| --- | --- |
| Astrocytes | GFAP, EAAT1, AQP4, LCN2, GJA1, SLC1A2, FGFR3, NKAIN4 |
| Endothelial cells | FLT1, CLDN5, VTN, ITM2A, VWF, FAM167B, BMX, CLEC1B |
| Excitatory neurons  Pericytes | SLC17A6, SLC17A7, NRGN, CAMK2A, SATB2, COL5A1, SDK2, NEFM |
| Inhibitory neurons | SLC32A1, GAD1, GAD2, TAC1, PENK, SST, NPY, MYBPC1, PVALB, GABBR2 |
| Microglia | IBA-1, P2RY12, CSF1R, CD74, C3, CST3, HEXB, C1QA, CX3CR1, AIF-1 |
| Oligodendrocytes | OLIG2, MBP, MOBP, PLP1, MOG, CLDN11, MYRF, GALC, ERMN, MAG |
| Oligodendrocyte precursor cells | VCAN, CSPG4, PDGFRA, SOX10, NEU4, PCDG15, GPR37L1, C1QL1, CDO1, EPN2 |
| Pericytes | AMBP, HIGD1B, COX4I2, AOC3, PDE5A, PTH1R, P2RY14, ABCC9, KCNJ8, CD248 |

**Supplementary Table S5**. The selection of differential gene expression analysis between different conditions (Condition 1 v.s. Condition 2) for diverse cell types in scREAD.

| Species | If in the same region | Condition 1 | Condition 2 |
| --- | --- | --- | --- |
| Human | Yes | Control | Disease |
| Mouse | Yes | Control | Disease |
| Human | Yes | Disease | Disease |
| Mouse | Yes | Disease | Disease |
| Human | No | Disease | Disease |
| Mouse | No | Disease | Disease |

*The comparisons are all in the same gender and age.

**Supplementary Table S6**. The computational tools used in scREAD.

| Tools | Source code | Version | Language |
| --- | --- | --- | --- |
| IRIS3 | https://github.com/OSU-BMBL/IRIS3 | v1.2.4 | R |
| Seurat | https://github.com/satijalab/seurat | v3.2 | R |
| Harmony | https://github.com/immunogenomics/harmony | v0.1 | R/Python |
| Polychrome | https://github.com/cran/Polychrome | v1.2.5 | R |
| Destiny | https://github.com/theislab/destiny/ | v3.2.0 | R |
| SCINA | https://github.com/jcao89757/SCINA | v1.0.0 | R |

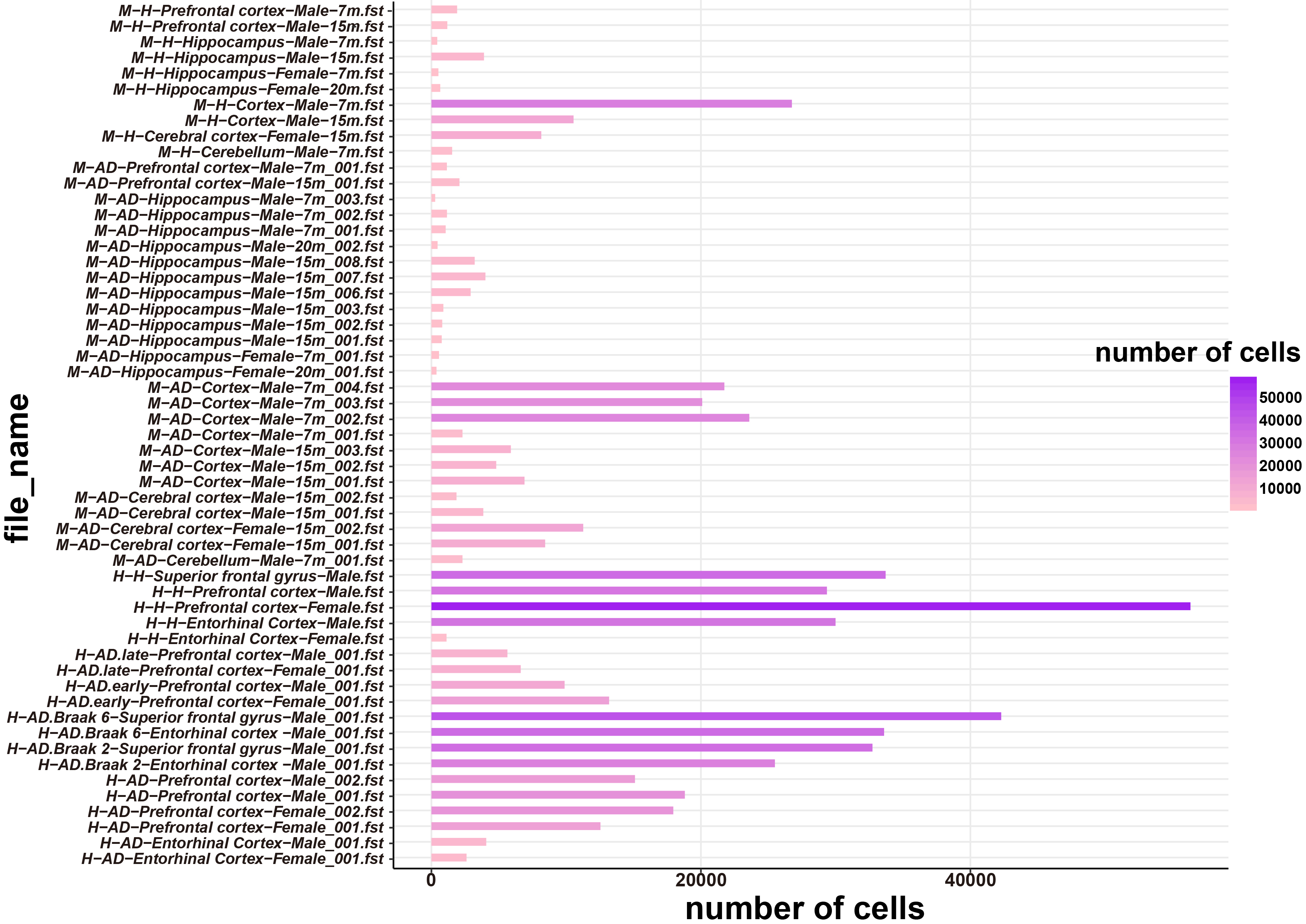

**Supplementary Figure S1. The number of cells in each of 55 files.** The x-axis represents the number of cells of each file, and the y-axis represents the file names of these 55 files. The color intensity of the bar stands for the number of cells, i.e. the darker of the color represents the more cell numbers in the corresponding file.

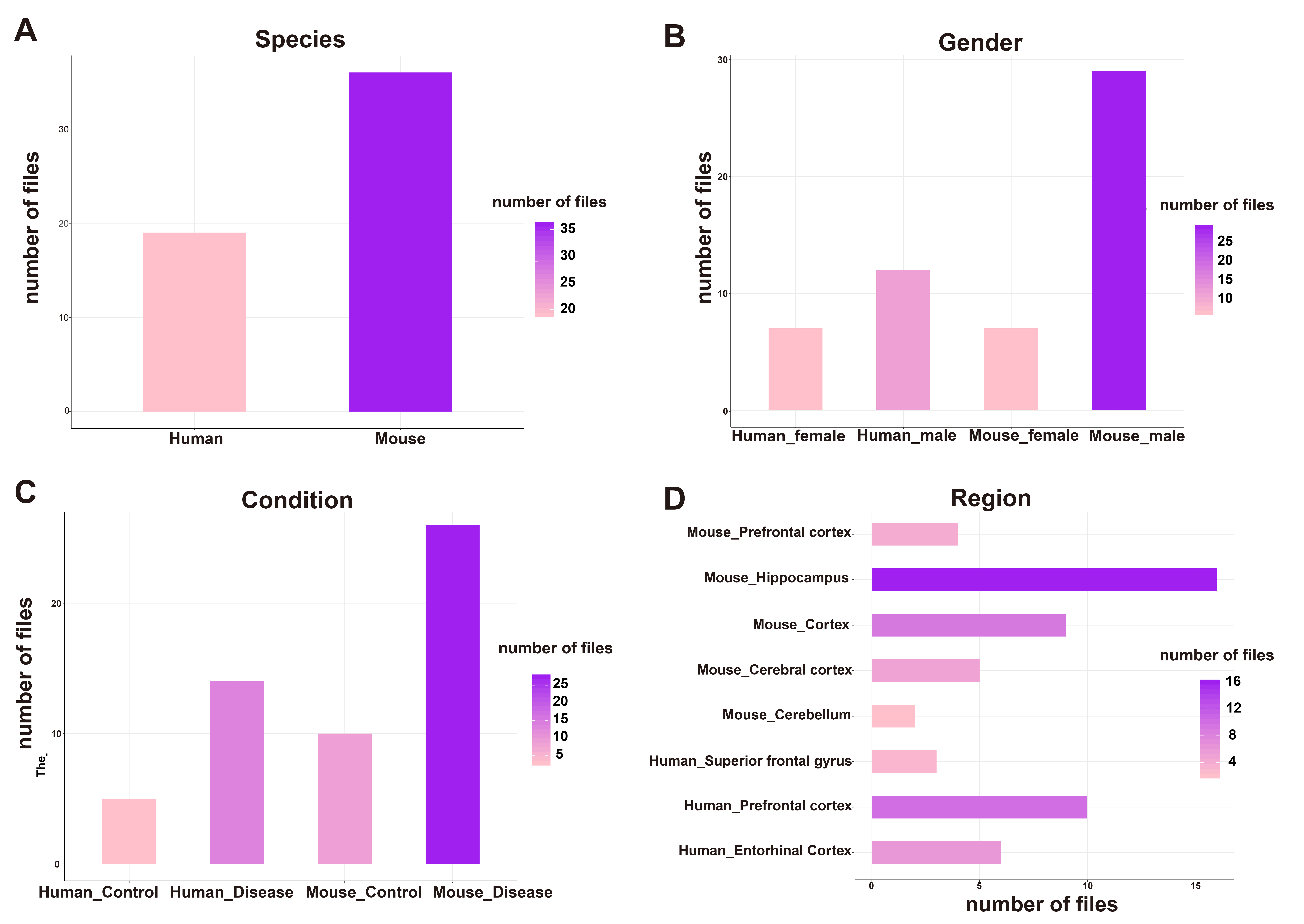

**Supplementary Figure S2. The distribution of the species, gender, condition, and brain region for 55 files.** For each panel in this figure, the color of the bar stands for the number of files, the darker the color is the more files in the corresponding factor.
